# The WD40 domain of autophagy protein ATG-16.2 is required for lysosomal regulation and longevity in germline-less *C. elegans*

**DOI:** 10.64898/2026.10.14.758959

**Authors:** Hiroshi Ebata, Sahiba K Dogra, Ling-Hsuan Sun, Debra Buggs, Saam Doroodian, Allison Linkous, Molly Hodul, Aimee W Kao, Malene Hansen

**Author notes:** Correspondence: M.H.

## Abstract

Autophagy is a conserved cellular recycling pathway linked to longevity and regulated by autophagy-related (ATG) protein. Many of these proteins serve a key function in ATG8 lipidation onto autophagosomal membranes, a step that precedes subsequent lysosomal degradation. Recently, mammalian ATG16L1, a member of ATG8 lipidation machinery, have been shown to also direct ATG8 lipidation onto other membranes via its C-terminal WD40 domain in acutely stressed cells. However, whether such non-canonical functions of the ATG16L1 WD40 domain influence longevity remain largely unknown. Here, we endogenously deleted the WD40 domain of *atg-16.2*, the *C. elegans* ortholog of *ATG16L1*, and examined autophagy, healthspan and longevity. The ATG-16.2 WD40 domain was dispensable for healthspan measures and for ATG8 lipidation in whole animals and neurons but was required for autophagic flux across all major tissues examined. Notably, the ATG-16.2 WD40 domain was also critical in limiting lysosomal membrane damage after acute heat stress. Because long-lived *C. elegans* mutants are typically heat resistant, we asked whether this lysosomal function supported their thermotolerance and found the WD40 domain to be selectively required for thermotolerance induced by germline removal, but not by reduced insulin/IGF-1 signaling or dietary restriction. In germline-less animals, loss of the ATG-16.2 WD40 domain reduced nuclear TFEB/HLH-30, reduced neuronal autophagic degradation, altered neuronal lysosomal pH, and sensitized animals to heat-induced lysosomal damage. Collectively, these findings establish the ATG16 WD40 domain as a regulator of autophagic degradation and lysosomal integrity *in vivo,* and show that this role is associated with longevity in a pathway-selective manner.

**Highlights:**

- The ATG-16.2 WD40 domain is dispensable for *C. elegans* healthspan but required for autophagic degradation *in vivo*
- The WD40 domain limits lysosomal membrane damage after acute heat stress *in vivo*
- The WD40 domain is selectively required for thermotolerance and longevity induced by germline removal (*glp-1/NotchR* mutation)
- Elevated neuronal lysosomal pH in germline-less animals requires the WD40 domain

## INTRODUCTION

Macroautophagy (hereafter autophagy) is a conserved cellular recycling process, by which cytosolic cargo is sequestered into double-membrane autophagosomes and delivered to lysosomes for degradation [1]. Autophagosome formation is carried out by a conserved set of autophagy-related (ATG) proteins acting in sequential steps [1], including conjugation of the ubiquitin-like protein ATG8 to the membrane lipid phosphatidylethanolamine. This lipidation marks the growing double-membrane phagophore that matures into the autophagosome, and is mediated by an E3-like complex of ATG5, ATG12 and ATG16 (ATG16L1 in humans and mammals) [1], within which ATG16 acts as the scaffold specifying the site of lipidation [1]. Autophagy dysfunction has been linked to aging and age-related diseases [2], whereas its activation can contribute to longevity [3], placing autophagy as a pro-longevity mechanism.

Beyond their canonical roles in autophagy, ATG proteins have recently been shown to also participate in additional, non-canonical pathways [4,5]. The best characterized form of such non-canonical pathways is the <u>c</u>onjugation of <u>A</u>TG8 proteins to <u>s</u>ingle <u>m</u>embranes (CASM) on endolysosomal compartments in response to diverse stresses, including pathogen invasion, lysosomotropic agents, and lysosomal membrane damage [4–6]. One of the key drivers of CASM is ATG16L1 [7], which, together with the same ATG12–ATG5 conjugate as in autophagy, directs ATG8 lipidation onto single membranes rather than onto forming double-membrane autophagosomes. What distinguishes CASM is the requirement for the C-terminal WD40 domain of ATG16L1, which directs ATG8 lipidation to endolysosomal membranes through its interaction with the V-ATPase [8]. Functionally, CASM has been proposed to act protectively: it promotes clearance of intracellular pathogens and supports lysosomal repair following membrane damage, and its loss is associated with infection and neurodegeneration [4–6]. Lysosomal dysfunction is a recognized feature of aging [9]: permeabilization of the lysosomal limiting membrane releases luminal hydrolases, abolishes lysosomal degradative capacity and, when extensive, triggers cell death. Yet, whether such non-canonical ATG functions including CASM contribute to organismal aging, however, remains largely unexamined.

Insights into how the WD40 domain functions have come primarily from mechanistic studies in mammalian cell cultures, in which lysosomal stress such as perturbed pH or ionic balance triggers WD40-dependent ATG8 lipidation on endolysosomal membranes [8,10,11]. In such lysosomal stress conditions, the WD40 domain is required for activating the transcription factor TFEB [12–14], a master regulator of lysosomal and autophagy gene expression [15], as well as recruiting lysosomal repair proteins [12,16–18]. These insights derive almost entirely from contexts using acute, non-physiological stress such as lysosomotropic agents in cultured cells, leaving the *in vivo* significance of the ATG16 WD40 domain poorly understood. A notable exception is the finding that mice expressing WD40-deficient ATG16L1 develop age-dependent neurodegeneration [19], implicating the WD40 domain in neuronal maintenance during aging. However, whether the ATG16 WD40 domain has a broader physiological role beyond a neuroprotective role during aging has yet to be investigated.

Given the non-canonical role of CASM at endolysosomal membranes, an important question is whether the WD40 domain also contributes to ATG16L1’s role in autophagy. The available *in vivo* evidence argues against such a contribution. Mice expressing WD40-deficient ATG16L1 showed no evident autophagy defect in kidney, liver, muscle or brain, based on steady-state LC3-I, LC3-II and p62/SQSTM1 levels measured by immunoblot of tissue lysates [19,20]. In *C. elegans*, a heat-shock-driven transgene lacking the WD40 domain of ATG16L1/ATG-16.2 rescues degradation of the autophagy receptor p62/SQST-1 in *atg-16.2* loss-of-function embryos [21]. Together these results led to the prevailing view that the domain is dispensable for autophagy. Such steady-state measurements, however, report marker abundance at a single time point and cannot distinguish increased autophagosome formation from impaired lysosomal degradation; resolving this requires measuring autophagic flux, in which marker levels are compared with and without lysosomal inhibition, or assessing lysosomal degradative capacity directly. Where these approaches have been applied in cultured cells, the results have differed. In several mammalian cell lines, WD40-deleted ATG16L1 supports autophagic flux, assessed by LC3-II and p62/SQSTM1 on immunoblot and LC3 puncta formation with and without a lysosomal inhibitor [19,20,22–24]. In contrast, loss of the WD40 domain delays ATG8 lipidation during starvation and the nutrient-sensing kinase mTOR measured as LC3-II time courses on immunoblot and suppresses basal turnover of long-lived proteins to the level of ATG16L1-null cells, defects attributed to weakened ATG16L1 binding to the early-acting autophagy proteins FIP200 and WIPI2 [25]. These differences may reflect cell type, stimulus or assay, and cultured cells cannot settle whether the WD40 domain is required for autophagy in a differentiated tissue, where autophagic demand and the lysosomal environment differ from those of proliferating cell lines. Whether the WD40 domain is required for autophagy in an intact animal, across tissues, has therefore not been tested.

Our previous *C. elegans* study implicated a link between a non-canonical role of the ATG-16.2 WD40 domain and autophagy [26]. *C. elegans* encodes two ATG16L1 orthologs, ATG-16.1 and ATG-16.2, each with a conserved C-terminal WD40 domain, of which ATG-16.2 has been shown to be the major contributor to autophagy in embryos [21]. We previously found that the ATG-16.2 WD40 domain is required for neuronal exopher formation that mediates the beneficial neuronal effects of inhibiting early-acting autophagy genes functioning until autophagosome formation [26]. However, that study relied on tissue-specific transgenes and did not address whether the endogenous ATG-16.2 WD40 domain contributes to organismal health and longevity. *C. elegans* is well-suited to address these questions: autophagic flux can be measured directly in intact tissues of this transparent nematode [27,28], and multiple conserved pathways and processes extend lifespan in an autophagy-dependent manner [2]. For example, knockdown of multiple, early-acting autophagy genes shortens the extended lifespan of animals with reduced insulin/IGF-1 signaling (IIS), of dietary-restricted animals (DR), and of germline-less animals [2]. Yet, whether WD40-mediated functions of ATG-16.2 contribute to these conserved longevity paradigms has never been addressed.

Here we asked what role the ATG-16.2 WD40 domain plays in the autophagy-lysosome pathway of an intact animal, and whether that role matters as the animal ages. We deleted the WD40 domain of *atg-16.2* from its endogenous locus by CRISPR-Cas9 and measured autophagy and healthspan in the intact animal. The domain proved dispensable for baseline healthspan and ATG8/LGG-1 lipidation at the whole-body and neuronal level yet required for lysosomal degradation in neurons: a requirement corroborated by impaired autophagic flux in neurons and in the other somatic tissues examined. Consistently, the domain was needed to limit lysosomal membrane damage after acute heat stress and, among longevity paradigms, was selectively required for the thermotolerance and lifespan extension induced by germline removal. In germline-less animals, its loss altered HLH-30/TFEB nuclear localization, neuronal autophagic degradation and neuronal lysosomal pH. These findings identify the ATG-16.2 WD40 domain as a paradigm-selective determinant of longevity and link it, *in vivo*, to lysosomal regulation.

## RESULTS

### *C. elegans* animals lacking the ATG-16.2 WD40 domain display normal healthspan

To date, analyses of the ATG-16.2 WD40 domain in *C. elegans*, by us and others, have relied on transgenic expression [21,26], leaving open whether the domain is required for organismal health and autophagy in an otherwise unmodified animal. To investigate the *in vivo* function of the ATG-16.2 WD40 domain, we used CRISPR-Cas9-mediated genome editing [29–31] and generated *atg-16.2(sq2)*, an endogenous allele lacking the C-terminal WD40 domain (ΔWD40, residues L224–R534) of ATG-16.2. To confirm that a stable protein was made without the WD40 domain, we generated mScarlet-tagged full-length and ΔWD40 alleles and examined protein levels by immunoblotting. Both full-length and truncated proteins were readily detected, with ΔWD40 migrating at the reduced size predicted for residues 1–223, and expression levels were comparable in animals expressing full-length ATG-16.2 and ΔWD40 (**Figure 1A, B**).

**Figure 1:**
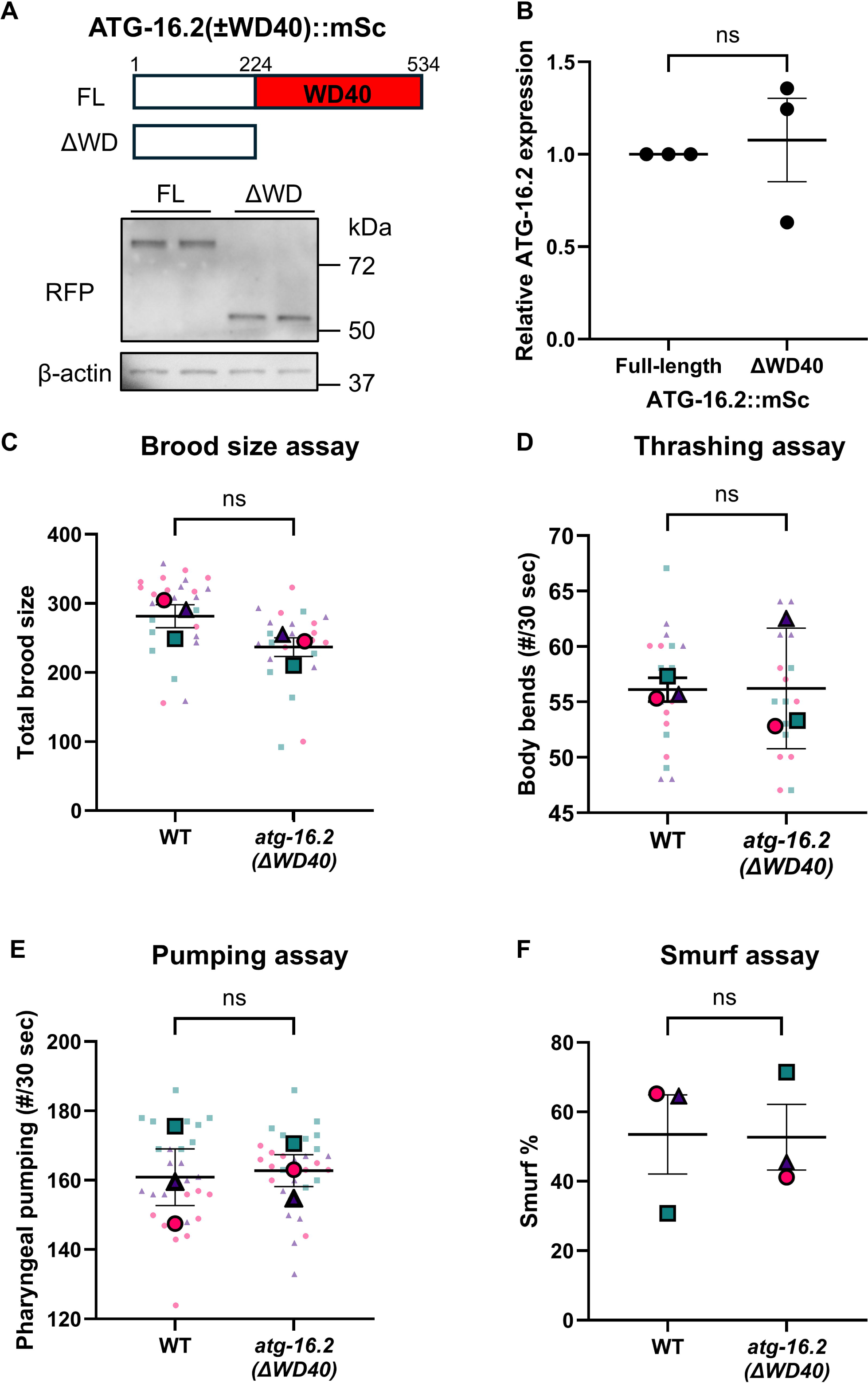
*atg-16.2(ΔWD40)* mutants do not show impairment in healthspan readouts. **A.** Representative immunoblot of ATG-16.2(±WD40)::mScarlet in an otherwise wild-type background on day 1 of adulthood, detected with an anti-RFP antibody; β-actin serves as a loading control. **B.** Quantification of ATG-16.2(±WD40)::mScarlet shown in (A) normalized to full-length strain within each replicate. Error bars show the mean ± standard error of the mean (SEM). n = 3. Welch’s t-test. Statistical tests were performed on raw measurements; values are displayed normalized to the WT control mean within each independent experiment for presentation. **C.** Total number of hatched eggs laid per animal was quantified in wild-type (WT, N2) and *atg-16.2(ΔWD40)* animals. Total brood size calculated from day 1 to day 5 of adulthood. Small dots represent data from each animal and large dots represent the means from independent experiments. Error bars show the mean ± standard error of the mean (SEM). n = 3. 25–26 animals. Welch’s t-test. **D.** WT and *atg-16.2(ΔWD40)* animals were assayed on day 1 of adulthood for thrashing ability in liquid M9 media. Small dots represent data from each animal and large dots represent the means from independent experiments. Error bars show the mean ± SEM. n = 3. 18 animals. Welch’s t-test. **E.** Number of pharyngeal pumps per minute were counted on day 1 of adulthood in WT and *atg-16.2(ΔWD40)* animals. Small dots represent data from each animal and large dots represent the means from independent experiments. Error bars show the mean ± SEM. n = 3. 30 animals. Welch’s t-test. **F.** Percent animals with blue dye in body cavity were quantified on day 10 of adulthood in WT and *atg-16.2(ΔWD40)* animals. Animals were soaked in blue food dye for 3 h. Animals containing blue dye leakages from the intestinal lumen into the body cavity were considered Smurf phenotype. Animals that presented no blue dye leakages from intestine to body cavity but contain dye in gonads were censored from the analysis. Error bars show the mean ± SEM. n = 3. 78–80 animals. Logistic regression.

Loss of autophagy function carries measurable costs to organismal fitness in *C. elegans*: core autophagy mutants more broadly show developmental issues and shortened lifespan [2]. While *atg-16.2* loss-of-function animals show reduced larval survival following starvation [21], whether ATG-16.2 and its WD40 domain are required for other aspects of adult fitness has not been examined. We therefore asked whether removing only the WD40 domain affects these animals’ intrinsic healthspan by measuring four fitness phenotypes [32]: brood size, thrashing, and pharyngeal pumping on day 1 of adulthood (the first day of egg-laying, when animals are reproductively active), and intestinal barrier integrity on day 10 of adulthood, assayed by leakage of a blue dye mixed with bacterial food (Smurf assay) [32,33]. Notably, *atg-16.2(ΔWD40)* animals were indistinguishable from wild type (WT) in all four measures (**Figure 1C–F**), in contrast to mutations in core autophagy genes [2].

### The ATG-16.2 WD40 domain is required for autophagic flux *in vivo*

Because ATG16 functions as part of the ATG8 lipidation machinery [1], we next asked whether the ATG-16.2 WD40 domain would be required for ATG8/LGG-1 lipidation *in vivo* under basal conditions (**Figure 2A**). ATG8/LGG-1 exists in an unlipidated form (LGG-1-I) and a lipidated, membrane-associated form (LGG-1-II), which can be resolved by immunoblot, and the LGG-1-II/LGG-1-I ratio is a standard measure of the extent of lipidation [27,34]. We quantified this ratio by immunoblot of endogenous ATG8/LGG-1 in whole-animal lysates on day 1 of adulthood. In *atg-5(lof*) animals, which display ATG8/LGG-1 lipidation defects in embryos [21], the LGG-1-II/LGG-1-I ratio was reduced, confirming ATG-5 is required for ATG8/LGG-1 lipidation (**Figure 2B**). In *atg-16.2(ΔWD40)* animals, by contrast, the ratio was not detectably different from WT (**Figure 2B**). This result suggests that the ATG-16.2 WD40 domain is dispensable for ATG8/LGG-1 lipidation *in vivo* at basal conditions, consistent with reports in mammalian cells that the domain is not required for ATG8 lipidation during autophagy [19,20,22–24].

**Figure 2:**
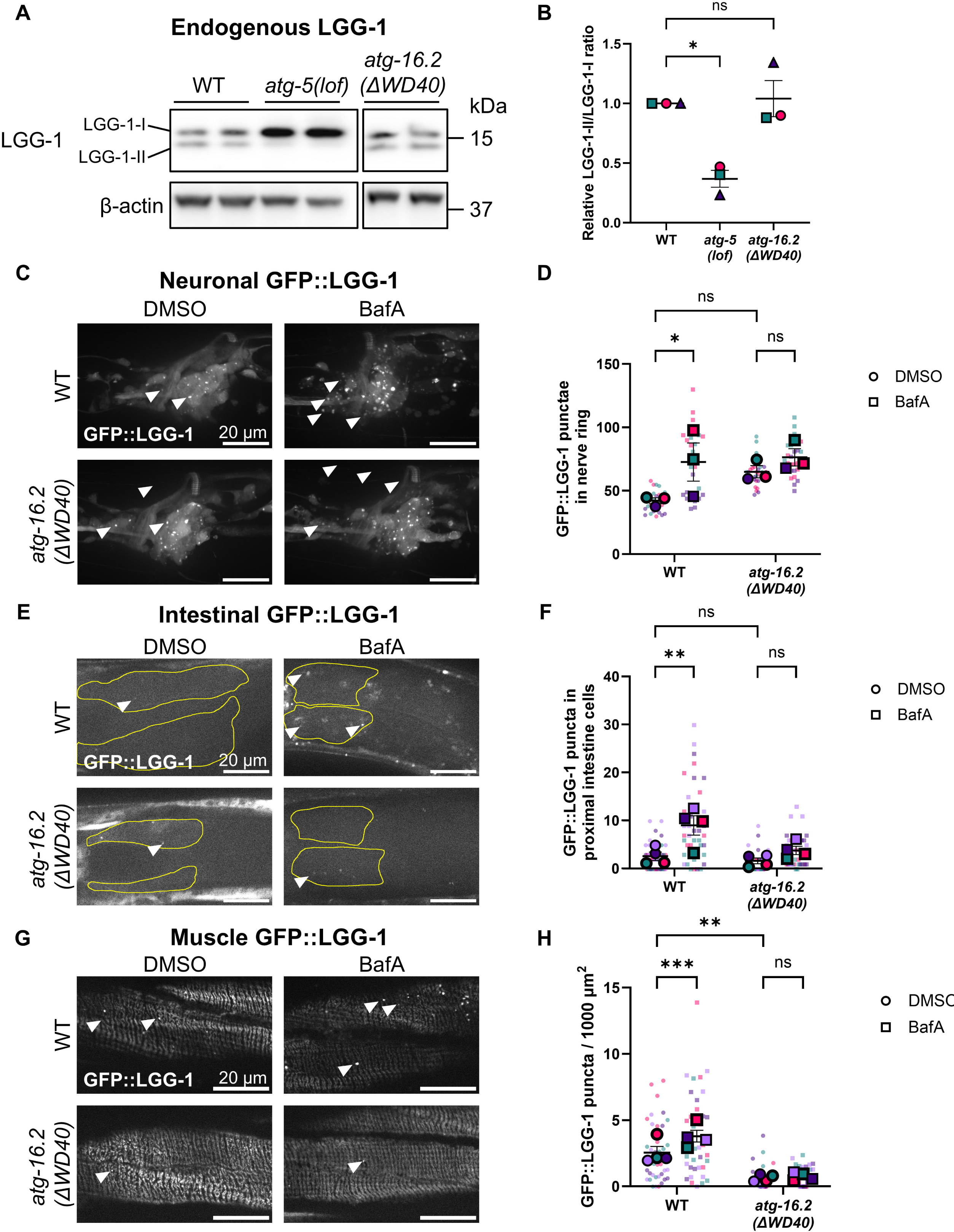
The ATG-16.2 WD40 domain is required for autophagic flux. **A.** Representative immunoblot of endogenous LGG-1 in WT, *atg-5(lof)* and *atg-16.2(ΔWD40)* animals on day 1 of adulthood, showing unlipidated LGG-1-I and lipidated LGG-1-II; β-actin serves as a loading control. **B.** Quantification of the LGG-1-II/LGG-1-I ratio shown in (A), normalized to WT within each replicate. Error bars show the mean ± SEM. n = 3. One-way ANOVA with Tukey’s multiple comparisons. *: p<0.05, Statistical tests were performed on log2 values of raw measurements; values are displayed normalized to the WT control mean within each independent experiment for presentation. **C.** Representative images of head regions in neuronal GFP::LGG-1/Atg8 reporter strains carrying WT or *atg-16.2(ΔWD40)* mutation on day 1 of adulthood with or without BafA treatment. Arrowheads indicate representative puncta. Scale bar: 20 μm. **D.** Quantification of GFP::LGG-1/Atg8 puncta in the nerve rings shown in (C). Small dots represent data from each animal and large dots represent the means from independent experiments. Error bars show the mean ± SEM. n = 3. 27–29 animals. Two-way ANOVA with Fisher’s LSD test . *: p<0.05. **E.** Representative images of proximal intestine in whole-body GFP::LGG-1/Atg8 reporter strains carrying WT or *atg-16.2(ΔWD40)* mutation on day 1 of adulthood with or without BafA treatment. Arrowheads indicate representative puncta. Scale bar: 20 μm. **F.** Quantification of GFP::LGG-1/Atg8 puncta in proximal intestinal cells shown in (E). Small dots represent data from each animal and large dots represent the means from independent experiments. Error bars show the mean ± SEM. n = 4. 39–41 animals. Two-way ANOVA with Fisher’s LSD test. **: p<0.01. **G.** Representative images of body wall muscles in whole-body GFP::LGG-1/Atg8 reporter strains carrying WT or *atg-16.2(ΔWD40)* mutation on day 1 of adulthood with or without BafA treatment. Arrowheads indicate representative puncta. Scale bar: 20 μm. **H.** Quantification of GFP::LGG-1/Atg8 puncta in body wall muscles shown in (G). Small dots represent data from each animal and large dots represent the means from independent experiments. Error bars show the mean ± SEM. n = 4. 39–41 animals. Two-way ANOVA with Fisher’s LSD test. *: p<0.05, **: p<0.01.

Because whole-animal lysates cannot resolve minor differences or changes confined to small tissues, we also asked whether the ATG-16.2 WD40 domain was required for autophagy *in vivo* tissue by tissue. To monitor autophagy, we used GFP::ATG8/LGG-1 reporters, which have been used to monitor pre-autophagosomal structures and autophagosomes, in three major somatic tissues (nerve-ring neurons, intestine and body-wall muscle) on day 1 of adulthood [27,35,36]. At baseline, *atg-16.2(ΔWD40)* mutants showed different profiles of GFP::LGG-1 puncta counts compared to WT in each tissue: in neurons, puncta counts were somewhat elevated relative to WT without reaching statistical significance (**Figure 2C, D**); in the intestine, counts were similar to WT (**Figure 2E, F**); and in body-wall muscle, counts were decreased (**Figure 2G, H**). These results suggest that the ATG-16.2 WD40 domain may regulate autophagosome formation differently in individual tissues.

To determine whether the observed GFP::LGG-1 puncta corresponded to lipidated, autophagosome-associated LGG-1, we created and examined a lipidation-deficient GFP::LGG-1(G116AG117A) reporter, which lacks the glycine residues exposed by the protease ATG-4 for lipidation [27,37]. Because lipidation recruits LGG-1 to membranes, this reporter is expected to form few puncta, and any formed puncta are considered as lipidation-independent. Consistent with a previous report using a neuronal GFP::LGG-1(G116A) reporter [38], GFP::LGG-1(G116AG117A) formed puncta at lower numbers than GFP::LGG-1 in WT neurons (∼73% reduction compared to GFP::LGG-1, **Figure S1A**). Critically, neuronal GFP::LGG-1(G116AG117A) puncta counts were comparable between WT and *atg-16.2(ΔWD40)* animals (**Figure S1A**), suggesting that loss of the WD40 domain does not alter this non-lipidated component. In the intestine and body-wall muscle, GFP::LGG-1(G116AG117A) puncta were rare in both genotypes (**Figure S1B, C**), indicating that GFP::LGG-1 puncta in these major tissues (**Figure 2E–H**) largely reflect lipidated structures. Together, these results show that the ATG-16.2 WD40 domain is required to maintain the steady-state abundance of lipidated GFP::LGG-1 structures, at least in body-wall muscle and possibly in neurons.

Because steady-state GFP::LGG-1 puncta numbers reflect the balance between autophagosome formation and their lysosomal degradation [27], these measurements alone cannot distinguish altered formation from altered turnover. We therefore quantified GFP::LGG-1 positive puncta in the presence of bafilomycin A1 (BafA), which inhibits the vacuolar H⁺-ATPase (V-ATPase) [39], the proton pump that acidifies the lysosomal lumen and thereby supports the activity of lysosomal hydrolases [40]. Inhibiting the V-ATPase blocks lysosomal degradation, so that autophagosomes accumulate when flux is intact but not when flux is already impaired (Zhang et al., 2015). In WT animals, BafA treatment (by top-coating method [28]) caused a robust increase in GFP::LGG-1 puncta in all tissues analyzed, confirming active autophagic flux in these tissues (**Figure 2C–H**), consistent with our previous findings [28]. In contrast, *atg-16.2(ΔWD40)* mutants showed no increase upon BafA treatment in any of the three tissues examined (**Figure 2C–H**). Considered together with the baseline GFP::LGG-1 puncta counts, these observations suggest that the ATG-16.2 WD40 domain may act at different steps of the pathway in different tissues. In body-wall muscle, baseline GFP::LGG-1 puncta were significantly reduced and did not increase upon BafA treatment (**Figure 2G, H**), the pattern expected if fewer autophagosomes were generated. In neurons and intestine, baseline counts were unchanged and likewise did not increase upon BafA treatment (**Figure 2C–F**). Altogether, our data indicate that autophagic flux was impaired in *atg-16.2(ΔWD40)* mutants.

As an additional measure of autophagy in these animals, we examined the level of p62/SQST-1, an autophagy receptor that is itself degraded by autophagy and is widely used as a readout of autophagy, with its accumulation indicating a block in lysosomal degradation [27]. Using an SQST-1::GFP reporter [26], GFP fluorescence intensity in the head region showed an increase in *atg-16.2(ΔWD40)* animals relative to WT on day 1 of adulthood (**Figure S2A, B**). We also examined polyglutamine (polyQ) aggregates, whose clearance depends on functional autophagy and whose accumulation reports a failure of aggregate degradation [41]. Neuronal polyQ aggregation (Q40::YFP) on day 7 of adulthood was increased in *atg-16.2(ΔWD40)* mutants compared to WT (**Figure S2C, D**). Similarly, in body-wall muscle, polyQ aggregation (Q35::YFP) on day 4 adulthood was increased in *atg-16.2(ΔWD40)* mutants compared to WT animals (**Figure S2E, F**). Collectively, the ATG-16.2 WD40 domain is required for autophagic flux in all three somatic tissues examined, with functional consequences for proteostasis, at least in neurons and body-wall muscle.

### The ATG-16.2 WD40 domain maintains neuronal lysosomal integrity under acute stress

Of the three tissues examined for autophagic flux, neurons were one of two in which polyQ aggregates accumulated in *atg-16.2(ΔWD40)* animals, and the tissue in which we previously implicated ATG-16.2 in organismal aging [26]. We therefore focused on neurons and used a neuron-specific GFP::LGG-1 reporter, which allows lipidation and degradation to be measured by immunoblotting as previously done with a ubiquitously expressed GFP::LGG-1 reporter [42]. Specifically, we measured GFP::LGG-1 lipidation as done above for whole animals and lysosomal degradation of GFP::LGG-1 as the amount of free GFP released upon lysosomal cleavage, relative to total GFP signal (**Figure 3A**). The GFP::LGG-1-II/GFP::LGG-1-I ratio in neurons did not show significant changes in *atg-16.2(ΔWD40)* mutants on day 3 of adulthood (**Figure 3B**), consistent with our observation that the loss of WD40 did not affect endogenous LGG-1 lipidation at whole-animal level (**Figure 2B**). Free GFP, by contrast, was reduced in *atg-16.2(ΔWD40)* animals relative to WT (**Figure 3C**). Because lipidation was unaffected, this reduction reflects less efficient lysosomal degradation rather than less lipidated GFP::LGG-1 and corroborates the absence of a BafA response in neurons (**Figure 2C, D**). The ATG-16.2 WD40 domain is therefore required for lysosomal degradation in neurons at a step distinct from ATG8/LGG-1 lipidation.

**Figure 3:**
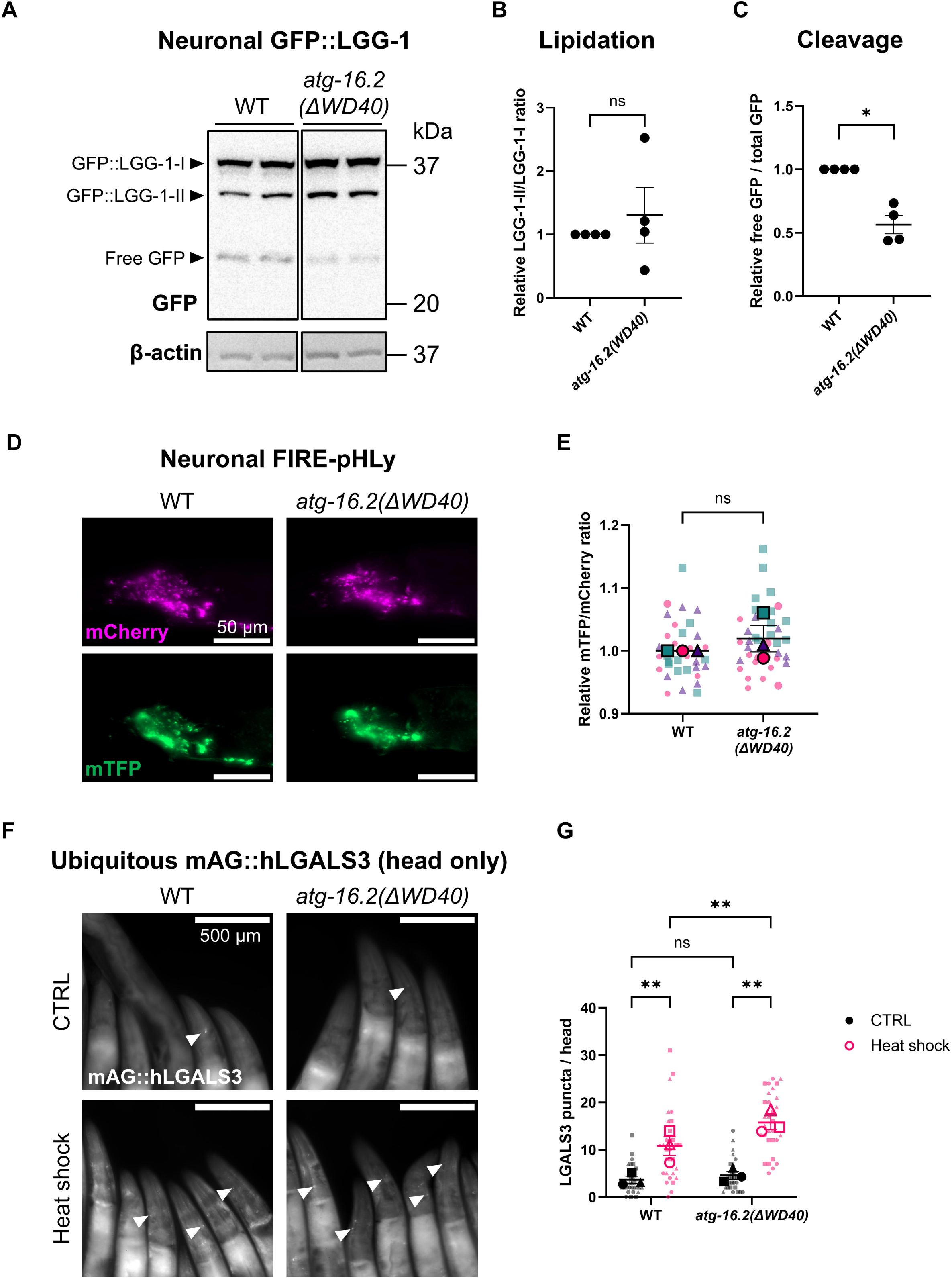
The WD40 domain of ATG-16.2 maintains lysosomal integrity under acute stress. **A.** Representative immunoblot of the neuronal GFP::LGG-1 cleavage assay in WT and *atg-16.2(ΔWD40)* animals on day 3 of adulthood, detected with an anti-GFP antibody, showing GFP::LGG-1-I, GFP::LGG-1-II and free GFP; β-actin serves as a loading control. **B.** Quantification of lipidation from (A), calculated as the GFP::LGG-1-II/GFP::LGG-1-I ratio and normalized to WT within each replicate. Error bars show the mean ± SEM. n = 4. Welch’s t-test. Statistical tests were performed on log2 values of raw measurements; values are displayed normalized to the WT control mean within each independent experiment for presentation. **C.** Quantification of degradation from (A), calculated as free GFP relative to total GFP signal and normalized to WT within each replicate. Error bars show the mean ± SEM. n = 4. Welch’s t-test. *: p<0.05. Statistical tests were performed on log2 values of raw measurements; values are displayed normalized to the WT control mean within each independent experiment for presentation. **D.** Representative images of neuronal FIRE-pHLy (mTFP::LMP-1::mCherry) reporter strains in WT and *atg-16.2(ΔWD40)* animals on day 3 of adulthood. Scale bar: 50 μm. **E.** Quantification of the mTFP/mCherry fluorescence intensity ratio shown in (D), normalized to WT within each replicate; a higher ratio indicates higher (less acidic) luminal pH. Small dots represent data from each animal and large dots represent the means from independent experiments. Error bars show the mean ± SEM. n = 3. 35–38 animals. Welch’s t-test. *: p<0.05. Statistical tests were performed on log2 values of raw measurements; values are displayed normalized to the WT control mean within each independent experiment for presentation. **F.** Representative images of head regions of mAG::hLGALS3 reporter animals in WT and *atg-16.2(ΔWD40)* backgrounds under control conditions (CTRL) and after heat shock (HS; 36 °C for 3 h followed by 3 h of recovery at 20 °C) on day 3 of adulthood. Arrowheads indicate representative puncta. Scale bar: 500 μm. **G.** Quantification of mAG::hLGALS3 puncta in the head region shown in (F). Small dots represent data from each animal and large dots represent the means from independent experiments. Error bars show the mean ± SEM. n = 3. 30 animals. Two-way ANOVA with Fisher’s LSD test. **: p<0.01.

The impaired GFP::LGG-1 degradation in neurons of *atg-16.2(ΔWD40)* mutants prompted us to analyze their lysosomal quality, a feature recently reported to be regulated by the ATG16L1 WD40 domain in mammalian cell cultures [6]. Thus, we focused on lysosomal luminal pH as a quantitative and physiologically central lysosomal parameter. To measure lysosomal pH, we used FIRE-pHLy, a ratiometric biosensor in which a higher mTFP/mCherry ratio reflects higher (less acidic) luminal pH [43], newly adapted for *C. elegans* and expressed under neuron-specific promoter. Using this new reporter, we found that neuronal lysosomal pH in *atg-16.2(ΔWD40)* animals was indistinguishable from WT (**Figure 3D, E**). The reduced autophagic degradation in the neurons of these animals (**Figure 3C**) is therefore not likely to arise from a change in steady-state lysosomal pH.

Recent cell culture studies show that the ATG16L1 WD40 domain is required for lysosomal stress response, including recruitment of lysosomal repair proteins, whose failure leads to loss of lysosomal integrity and accumulation of ruptured lysosomes [6]. We therefore asked whether lysosomal integrity is compromised under conditions that challenge the lysosome directly. Lysosomal membrane rupture exposes glycoconjugates on the luminal side of lysosomal membrane proteins, which is detected by galectin proteins such as galectin-3 (LGALS3) [44]. Thus, LGALS3 can visualize damaged lysosomes as LGALS3 puncta. In *C. elegans*, a previous study established a human LGALS3 reporter tagged with mAzamiGreen (mAG) for lysosomal damage assay driven by a ubiquitous promoter [45]. Using this reporter, we counted mAG::hLGALS3 puncta in the head region (**Figure 3F**), which contains the nerve ring, and across the whole body (**Figure S3A**) after 36°C heat shock, which has been shown to be sufficient to induce lysosomal damage in the *C. elegans* hypodermis [46]. Under basal conditions, mAG::hLGALS3 puncta were comparable between WT and *atg-16.2(ΔWD40)* animals both in head regions as well as whole body (**Figure 3F, G**; Figure S3A, B). However, following heat shock, mAG::hLGALS3 puncta increased in both strains, and were more numerous in *atg-16.2(ΔWD40)* animals than in WT (**Figure 3G**; **Figure S3B**). This is consistent with the ATG-16.2 WD40 domain limiting lysosomal membrane damage following acute stress, consistent with the domain’s role described in mammalian cells [6]. Because the difference between genotypes was larger across the whole body than in the head region, this requirement does not appear to be restricted to neurons. Since the increased lysosomal membrane damage in *atg-16.2(ΔWD40)* animals after heat shock was detected both in head regions and across the whole body, this requirement of the ATG-16.2 WD40 domain for limiting lysosomal damage does not appear to be confined to neurons.

### The ATG-16.2 WD40 domain is differentially required across *C. elegans* longevity paradigms for thermotolerance

A general feature of *C. elegans* longevity mutants is enhanced resistance to acute stress, including thermotolerance [2,47]. Because our data indicated that the ATG-16.2 WD40 domain was required for autophagic flux across tissues (**Figure 2C, D**; Figure S1) and limited lysosomal damage following heat stress (**Figure 3D**), we asked whether the WD40 domain was similarly required for thermotolerance conferred by multiple conserved longevity pathways. Specifically, we tested three well-characterized longevity mutants by crossing *atg-16.2(ΔWD40)* allele into these mutants: the insulin/IGF-1 receptor *daf-2(e1370)* (reduced insulin/IGF-1 signaling) [35,48], the acetylcholine receptor *eat-2(ad1116)* (dietary restriction) [49,50], and the Notch receptor *glp-1(e2144)* (germline removal) [51,52].

To assess thermotolerance, we exposed animals to 36°C on day 3 of adulthood and scored survival after recovery. Introducing the *atg-16.2(ΔWD40)* allele did not detectably change the thermotolerance of *eat-2(ad1116)* mutants (**Figure 4A**) or of *daf-2(e1370)* mutants (**Figure 4B**), indicating that the ATG-16.2 WD40 domain is not required for the stress resistance induced by reduced insulin/IGF-1 signaling or by dietary restriction. In contrast, germline-deficient Notch receptor *glp-1(e2144)* mutants carrying *atg-16.2(ΔWD40)* showed reduced thermotolerance compared to *glp-1(e2144)* alone (**Figure 4C**), suggesting that the WD40 domain of ATG-16.2 is required for germline-removal induced stress resilience. Importantly, ATG-16.2(ΔWD40)::mScarlet was expressed at a comparable level to full-length ATG-16.2::mScarlet in *glp-1(e2144)* background (**Figure 4D, E**), suggesting that the decrease in thermotolerance in *atg-16.2(ΔWD40); glp-1(e2144)* mutants is not due to unstable ATG-16.2(ΔWD40). The ATG-16.2 WD40 domain is therefore selectively required, among these three longevity paradigms, for the thermotolerance of germline-less animals.

**Figure 4:**
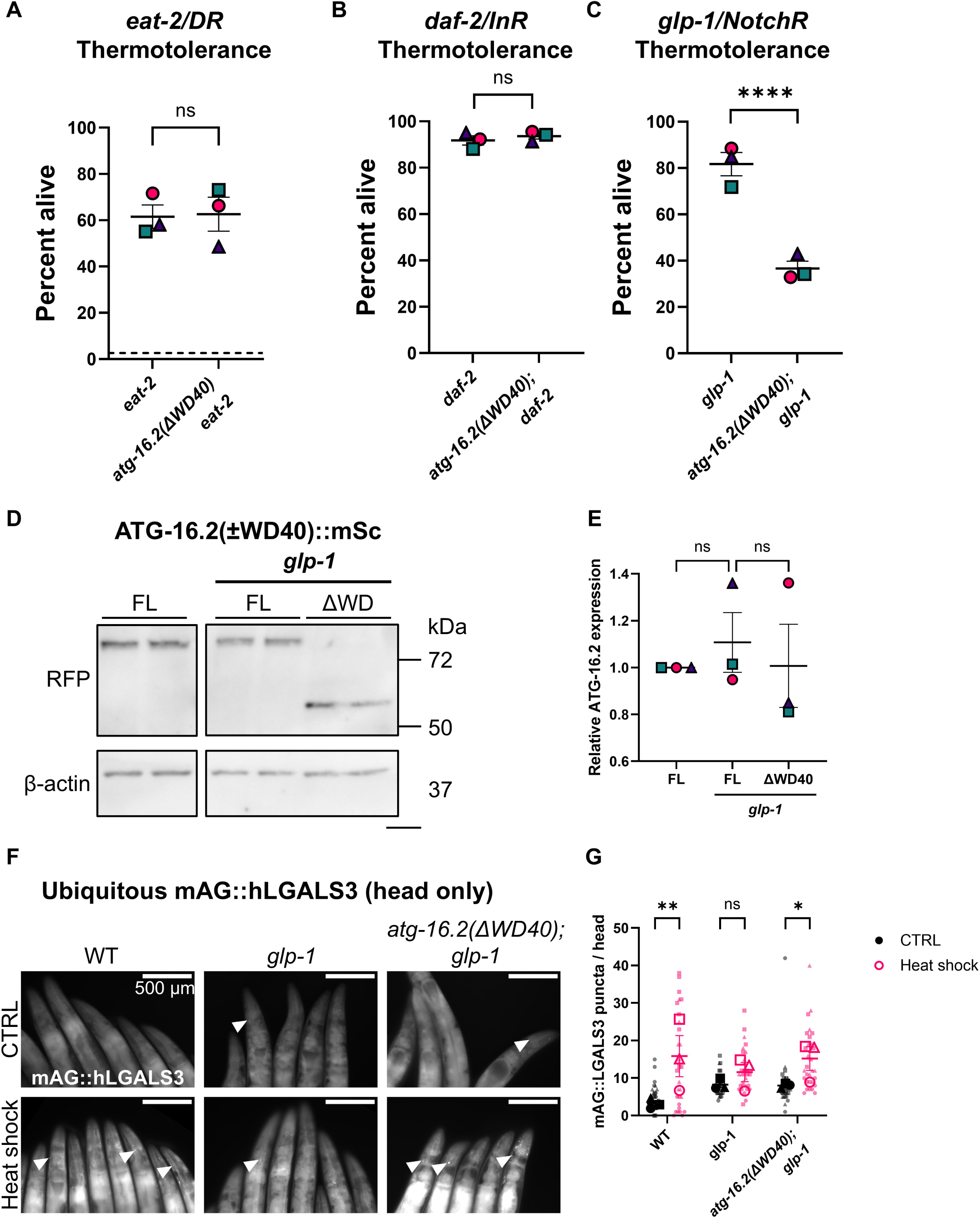
The WD40 domain of ATG-16.2 is required for thermotolerance conferred by germline removal. **A.** Thermotolerance of *eat-2(ad1116)* mutants carrying WT or *atg-16.2(ΔWD40)* mutation on day 3 of adulthood. Heat shock was introduced at 36 °C for 5 hours, and survival was scored ∼24 h later. Error bars show the mean ± SEM. A dot line represents viability of wild-type animals after the same heat shock regime. N = 3. 300–311 animals. Logistic regression. ****: p<0.0001. **B.** Thermotolerance of *daf-2(e1370)* mutants carrying WT or *atg-16.2(ΔWD40)* mutation on day 3 of adulthood. Heat shock was introduced at 36 °C for 9 hours, and survival was scored ∼48 h later. Error bars show the mean ± SEM. n = 3. 174–215 animals. Logistic regression. **C.** Thermotolerance of *glp-1(e2144)* mutants carrying WT or *atg-16.2(ΔWD40)* mutation on day 3 of adulthood. Heat shock was introduced at 36 °C for 6 hours, and survival was scored ∼24 h later. Error bars show the mean ± SEM. n = 3. 222–235 animals. Logistic regression. ****: p<0.0001. **D.** Representative immunoblot of ATG-16.2(±WD40)::mScarlet in the *glp-1(e2144)* background, detected with an anti-RFP antibody; β-actin serves as a loading control. **E.** Quantification of ATG-16.2(±WD40)::mScarlet levels shown in (D), normalized to full-length ATG-16.2::mScarlet control within each replicate. Error bars show the mean ± SEM. n = 3. One-way ANOVA with Tukey’s multiple comparisons. Statistical tests were performed on log2 values of raw measurements; values are displayed normalized to the WT control mean within each independent experiment for presentation. **F.** Representative images of mAG::hLGALS3 reporter animals in WT, *glp-1(e2144)* and *atg-16.2(ΔWD40); glp-1(e2144)* backgrounds under control conditions (CTRL) and after heat shock (HS; 36 °C for 3 h followed by 3 h of recovery at 20 °C) on day 3 of adulthood. Arrowheads indicate representative puncta. Scale bar: 500 μm. **G.** Quantification of mAG::hLGALS3 puncta in the head region shown in (F). Small dots represent data from each animal and large dots represent the means from independent experiments. Error bars show the mean ± SEM. n = 3. 30 animals. Two-way ANOVA with Tukey’s multiple comparisons. *: p<0.05, **: p<0.01.

The selective requirement in germline-less animals for thermotolerance raised the question of whether the lysosomal damage phenotype we observed in *atg-16.2(ΔWD40)* animals would also be observed in germline-less animals. We therefore measured mAG::LGALS3 puncta in the head region and across the whole body in WT, *glp-1(e2144)* and *atg-16.2(ΔWD40); glp-1(e2144)* animals before and after heat shock (**Figure 4F; Figure S3C**). Following heat shock, mAG::LGALS3 puncta increased in WT animals but not in *glp-1(e2144)* mutants both in the head region (**Figure 4F, G**) and across the whole body (**Figure S3C, D**), suggesting that germline removal induces protection against heat-induced lysosomal membrane damage. Interestingly, in the double mutants, that protection was lost: mAG::LGALS3 puncta increased after the heat shock (**Figure 4G; Figure S3D**). These results indicate that germline removal protects lysosomes from acute heat stress, and this protection requires the ATG-16.2 WD40 domain. Together, these results suggest that the ATG-16.2 WD40 domain is required both for enhanced thermotolerance in *glp-1/NotchR* mutants and for their protection against heat induced lysosomal damage.

### The ATG-16.2 WD40 domain is linked to lysosomal regulation in *glp-1/NotchR* mutants

Thermotolerance is considered as a robust correlate of longevity [47]. Since we observed the requirement of the ATG-16.2 WD40 domain for thermotolerance in germline-less animals, we asked whether the ATG-16.2 WD40 domain is also important for longevity of germline-less animals. As previously reported [53], *glp-1(e2144)* animals lived substantially longer than WT (**Figure 5A**, **Table 1**). Importantly, this extension was reduced in *atg-16.2(ΔWD40); glp-1(e2144)* double mutants (**Figure 5A**, **Table 1**). The ATG-16.2 WD40 domain is therefore required for the full lifespan extension induced by germline removal.

**Figure 5:**
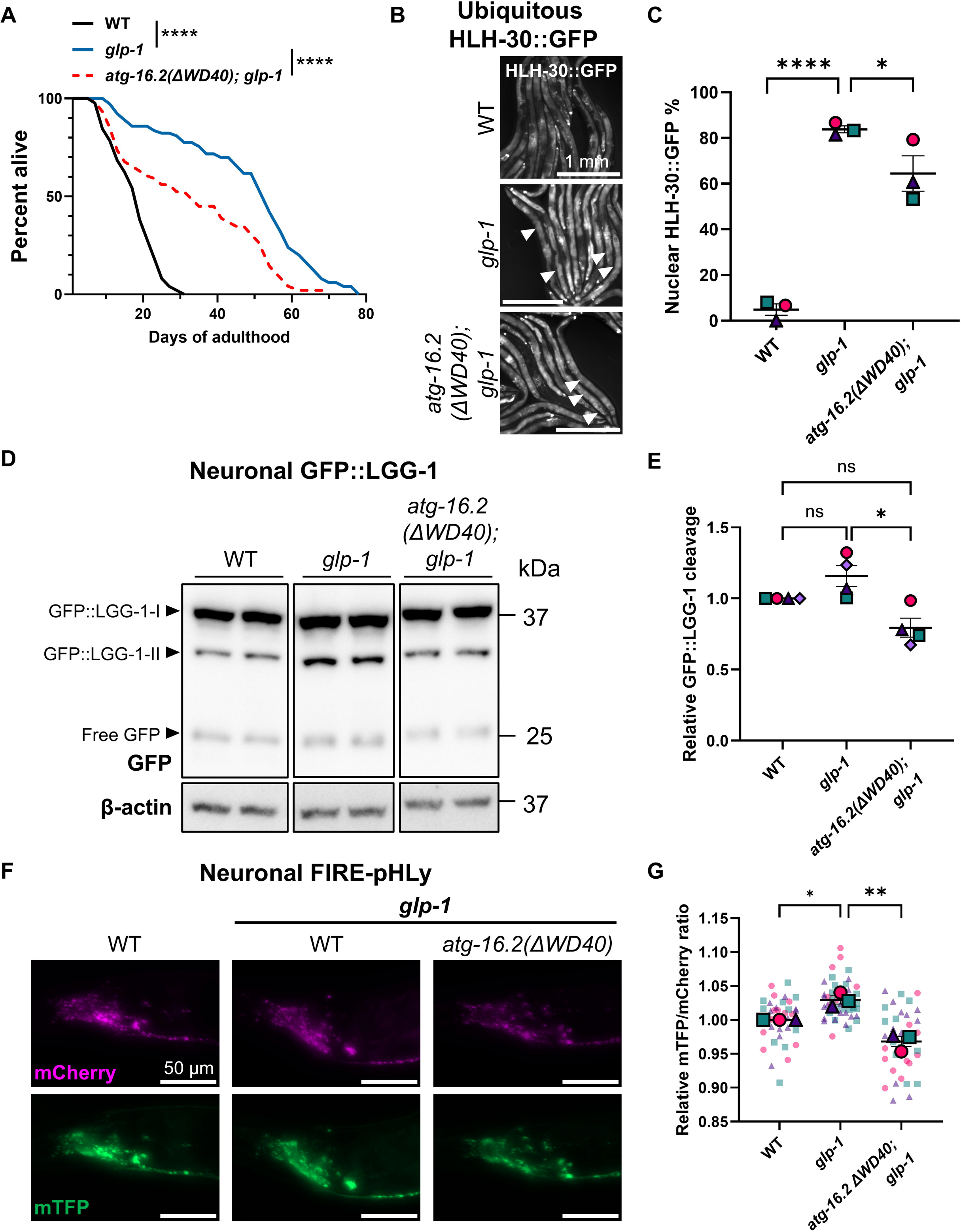
Loss of the ATG-16.2 WD40 domain abolishes longevity and the elevated neuronal lysosomal pH characteristic of *glp-1* mutants. **A.** Lifespan of WT, *glp-1(e2144)* and *atg-16.2(ΔWD40);glp-1(e2144)* animals. n = 5. 120 animals. Log-rank test. ****: p<0.0001. See Table 1 for a summary of all lifespan experiments. **B.** Representative images of HLH-30::GFP reporter animals in WT, *glp-1(e2144)* and *atg-16.2(ΔWD40); glp-1(e2144)* backgrounds on day 3 of adulthood. Arrowheads indicate representative nuclear localization. Scale bar: 1 mm. **C.** HLH-30::GFP nuclear localization, expressed as the percentage of animals scored as nuclear on day 3 of adulthood. Error bars show the mean ± SEM. n = 3. 81–87 animals. Logistic regression with Tukey’s multiple comparisons. *: p<0.05, ****: p<0.0001. **C.** Representative immunoblot of the neuronal GFP::LGG-1 cleavage assay in WT, *glp-1(e2144)* and *atg-16.2(ΔWD40); glp-1(e2144)* animals on day 3 of adulthood, detected with an anti-GFP antibody, showing GFP::LGG-1-I, GFP::LGG-1-II and free GFP; β-actin serves as a loading control. **D.** Quantification of free GFP relative to total GFP signal shown in (D), normalized to WT within each replicate. Error bars show the mean ± SEM. n = 4. One-way ANOVA with Tukey’s multiple comparisons. *: p<0.05. Statistical tests were performed on log2 values of raw measurements; values are displayed normalized to the WT control mean within each independent experiment for presentation. **E.** Representative images of neuronal FIRE-pHLy reporter strains in WT, *glp-1(e2144)* and *atg-16.2(ΔWD40); glp-1(e2144)* backgrounds on day 3 of adulthood. Scale bar: 50 μm. **F.** Quantification of mTFP/mCherry fluorescence intensity ratio shown in (F), normalized to WT within each replicate.. Small dots represent data from each animal and large dots represent the means from independent experiments. Error bars show the mean ± SEM. n = 3. 35–48 animals. One-way ANOVA with Tukey’s multiple comparisons. *: p<0.05, **: p<0.01. Statistical tests were performed on log2 values of raw measurements; values are displayed normalized to the WT control mean within each independent experiment for presentation.

**Table 1:** Summary of lifespan experiments of comparing wild-type, *glp-1* and *atg-16.2(sq2); glp-1*.

| Exp # | Strain Name | Strain Genotype | MLS (Days) | % MLS change | P value | N animals/ total |
| --- | --- | --- | --- | --- | --- | --- |
| 1 | N2 | wild type (WT) | 18.8 | -60 | <0.0001 | 67/120 |
|  | CF1903 | *glp-1(e2144)* | 47.5 |  |  | 45/120 |
|  | MAH1355 | *atg-16.2(sq2); glp-1(e2144)* | 30.3 | -36 | <0.0001 | 74/120 |
| 2 | N2 | WT | 17.9 | -63 | <0.0001 | 51/120 |
|  | CF1903 | *glp-1(e2144)* | 48.2 |  |  | 59/120 |
|  | MAH1355 | *atg-16.2(sq2); glp-1(e2144)* | 33.5 | -30 | <0.0001 | 68/120 |
| 3 | N2 | WT | 18.3 | -54 | <0.0001 | 88/120 |
|  | CF1903 | *glp-1(e2144)* | 40 |  |  | 66/120 |
|  | MAH1355 | *atg-16.2(sq2); glp-1(e2144)* | 22.4 | -44 | <0.0001 | 83/120 |
| 4 | N2 | WT | 18.6 | -57 | <0.0001 | 83/120 |
|  | CF1903 | *glp-1(e2144)* | 43.3 |  |  | 88/120 |
|  | MAH1355 | *atg-16.2(sq2); glp-1(e2144)* | 30.7 | -29 | <0.0001 | 66/120 |
| 5 | CF1903 | *glp-1(e2144)* | 47.4 |  |  | 52/120 |
|  | MAH1355 | *atg-16.2(sq2); glp-1(e2144)* | 35.8 | -24 | <0.0001 | 54/120 |

Longevity in germline-less animals also depends on HLH-30, the ortholog of the transcription factor TFEB, which accumulates in the nucleus of *glp-1(e2144)* animals under basal conditions and is required for their lifespan extension [54]. TFEB is a central regulator of lysosomal and autophagy gene expression [15], and the ATG16L1 WD40 domain is required for TFEB activation under lysosomal stress in mammalian cells [12,13], making TFEB/HLH-30 a candidate point of convergence. We therefore scored nuclear localization of HLH-30::GFP in intestinal cells similar to what we did previously [54] on day 3 of adulthood (**Figure 5B**). As reported, *glp-1(e2144)* animals showed increased nuclear HLH-30::GFP relative to WT (**Figure 5C**). This increase was partially but significantly reduced in *atg-16.2(ΔWD40); glp-1(e2144)* double mutants (**Figure 5C**), indicating that the ATG-16.2 WD40 domain is at least partially required for full HLH-30 nuclear localization in germline-less animals, and, notably, in the absence of any exogenous lysosomal stress.

Finally, we returned to the neuron-specific GFP::LGG-1 cleavage assay, which separates lipidation from degradation within a single tissue, to ask whether the WD40 domain is required for lysosomal function in *glp-1(e2144)* mutants under basal conditions. Free GFP was reduced in *atg-16.2(ΔWD40); glp-1(e2144)* animals compared to *glp-1(e2144)* animals (**Figure 5D, E**), indicating reduced lysosomal degradation in these mutants. GFP::LGG-1 lipidation in neurons assessed by GFP::LGG-1-II/GFP::LGG-1-I ratio was not altered by *atg-16.2(ΔWD40)* mutation in *glp-1(e2144)* animals (**Figure S4**), and thus, the impaired degradation in neurons is independent of GFP::LGG-1 lipidation. Next, we asked whether the ATG-16.2 Wd40 domain is required for lysosomal properties of neurons in germline-less animals, using neuron-specific FIRE-pHLy reporter. We found that lysosomal pH in neurons of *glp-1(e2144)* animals was elevated relative to WT (**Figure 5F, G**). This was unexpected, since lysosomal hydrolases require an acidic lumen {Maxson, 2014 #324}, and impaired lysosomal acidification is itself a reported feature of aging [9]; a higher luminal pH is therefore not the obvious correlate of a long-lived state. In *atg-16.2(ΔWD40); glp-1(e2144)* animals, this elevation of neuronal lysosomal pH in germline-less animals was abolished, returning pH to a level comparable to WT (**Figure 5G**). Taken together, we suggest that germline removal increases lysosomal pH in neurons, whereas loss of the ATG-16.2 WD40 domain acidifies lysosomes, indicating that the WD40 domain restrains lysosomal acidification in *glp-1/NotchR* animals. To our knowledge, these findings provide the first *in vivo* evidence that ATG16 WD40 function regulates lysosomal pH. Together, these associations place neuronal lysosomal function as a potential route, by which the ATG-16.2 WD40 domain contributes to thermotolerance and longevity induced by germline removal.

## DISCUSSION

Here, we found that an endogenous deletion of the ATG-16.2 WD40 domain impaired autophagic degradation across adult tissues of *C. elegans*. Surprisingly, despite this defect, healthspan was unaffected in the WD40-deficient mutants. Instead, further analyses revealed that the domain was required to limit lysosomal membrane damage after heat shock. Notably, the WD40 deletion reduced both the thermotolerance and the lifespan extension of germline-less *glp-1/NotchR* animals, but not the thermotolerance of *daf-2/InR* or *eat-2/DR* animals, highlighting the context-dependent requirement for the ATG-16.2 WD40 domain in organismal stress resilience. In *glp-1/NotchR* animals, the deletion of the ATG-16.2 WD40 domain also altered several lysosomal properties, including luminal pH in neurons. Together, these results implicate lysosomal function in how the ATG-16.2 WD40 domain contributes to longevity *in vivo*.

We observed the ATG-16.2 WD40 domain to be required for autophagic degradation across adult *C. elegans* tissues *in vivo*. Autophagic flux was impaired in neurons, intestine, and body-wall muscle of WD40-deleted animals, p62/SQST-1 increased in the head region, and polyQ aggregates accumulated in neurons and body-wall muscle. Furthermore, in neurons, degradation of GFP::ATG8/LGG-1 was impaired while its lipidation was unchanged, indicating that loss of the ATG-16.2 WD40 domain may affect a step distinct from ATG8/LGG-1 lipidation in autophagy. The affected step may differ between tissues: in body-wall muscle, basal GFP::LGG-1 puncta were reduced rather than increased, favoring a defect in autophagosome formation over one in lysosomal degradation. ATG8/LGG-1 lipidation was not measured in muscle, leaving open the possibility that the domain contributes to lipidation in this tissue. Previous studies examining the role of the ATG16 WD40 domain in autophagy reached differing conclusions. Studies in cultured mammalian cells and in mouse tissues found no autophagy defect when assessed by steady-state markers [19,20,22,24]. In *C. elegans* embryos, an overexpressed WD40-truncated ATG-16.2 transgene restored degradation of p62/SQST-1 in *atg-16.2* loss-of-function mutants, concluding that the domain is dispensable for autophagy, at least during *C. elegans* development [21]. In contrast, more recent work found that WD40 deletion weakens ATG16L1 binding to FIP200 and WIPI2 and delays LC3 lipidation in HeLa cells [25]. Our data add a third possibility: a requirement of the ATG-16.2 WD40 domain for autophagic degradation that is not explained by its effect on bulk ATG8/LGG-1 lipidation, at least in neurons. It is worth noting that our lipidation measurements cannot distinguish ATG8/LGG-1 lipidation onto autophagosomes from lipidation onto other membranes. Thus, we cannot exclude the possibility that target membrane of ATG8/LGG-1 lipidation is altered in *atg-16.2(ΔWD40)* mutants. Whether such conjugation at single membranes (akin to CASM) occurs in adult *C. elegans* at all, and whether the ATG-16.2 WD40 domain supports it, remain to be tested. The domain may alternatively act through an activity that does not involve conjugation or may function differently in *C. elegan*s and in mammalian cells. In each case, our data suggest that the ATG-16.2 WD40 domain is required for autophagy *in vivo* through a contribution that its established role in ATG8/LGG-1 lipidation may not account for.

The reduced degradation of GFP::LGG-1 in neurons without a lipidation defect in *atg-16.2(ΔWD40)* mutants pointed to potential impairment in lysosome quality and prompted us to examine lysosomal properties. Steady-state lysosomal pH in *atg-16.2(ΔWD40)* mutants’ neurons was indistinguishable from wild type, indicating that the degradation defect does not result from a failure in lysosomal acidification. Interestingly, we observed that the WD40 domain was required for lysosomal integrity after heat shock, both in the head region and across the whole body. Because the lysosomal damage phenotype was detectable across the whole body, the requirement of the ATG-16.2 WD40 domain seems not confined to neurons. Our assay counted damaged lysosomes at a single time point, which does not distinguish increased susceptibility to damage from impaired repair or clearance; a recovery time course of a lysosomal damage marker after heat shock will be needed to separate these possibilities for future investigation. The requirement of the ATG-16.2 WD40 domain for lysosomal damages responses is interesting in light of the reported role of the ATG16L1 WD40 domain in recruiting lysosomal repair machinery in mammalian cells [6]. Examining ATG-16.2 recruitment to lysosomes after heat shock would be needed to test whether a comparable relationship also exists *in vivo*.

Autophagic/lysosomal defects caused by loss of the ATG-16.2 WD40 domain were not accompanied by a corresponding fitness cost: *atg-16.2(ΔWD40)* animals showed normal brood size, thrashing, pharyngeal pumping, and intestinal barrier integrity. This contrasts to the reduced starvation survival reported for *atg-16.2* loss-of-function animals [21] as well as known developmental/healthspan defects in core autophagy mutants [2]. Our observations parallel the *Atg16l1^E230^*mouse, which lacks the WD domains at the endogenous locus [20]. These animals are born at Mendelian frequency, grow at the same rate as littermate controls, are fertile with normal litter sizes, and live at least 24 months [20]. Liver, kidney, brain and muscle show organ weights comparable to controls relative to body weight and no tissue damage at 2–3 months, with serum liver enzymes and inflammatory cytokines likewise unaffected at 2–6 months [20]. Yet, these mice are severely susceptible to viral, bacterial and parasite infection [55]. Therefore, loss of the WD domain leaves both the mouse and *C. elegans* essentially unimpaired under unstressed conditions. Several explanations for the absence of a fitness cost in *atg-16.2(ΔWD40) C. elegans* animals are possible. The truncated ATG-16.2 protein produced in the *atg-16.2(ΔWD40)* animals is expressed at levels comparable to full-length and may therefore support sufficient autophagy for normal development and fitness under normal conditions. *C. elegans* also encodes another ATG16L1 ortholog, ATG-16.1, which carries its own C-terminal WD40 domain and acts partially redundantly with ATG-16.2 in autophagy [21]. We did not examine *atg-16.1* here, and compensation by this paralog [21] could mask phenotypes in the assays we used; it may also explain why the effects we do observe, such as the reduction in lifespan and in HLH-30/TFEB nuclear localization in *glp-1/NotchR* mutants, are partial rather than complete. Establishing whether ATG-16.1 contributes, and whether its WD40 domain is functionally equivalent to that of ATG-16.2, will be of interest to address in future experiments.

A striking result was that the ATG-16.2 WD40 domain is required for *glp-1/NotchR* mediated thermotolerance but dispensable for two other longevity paradigms, i.e., *daf-2/InR* and *eat-2/DR* mutations. This selectivity is notable because all three paradigms have been reported to require multiple autophagy genes [2], albeit with different tissue requirement [2]. We did not measure autophagic flux in longevity backgrounds, and thus, whether the WD40 deletion could impair autophagy in these animals remains untested. Alternatively, *daf-2/InR* and *eat-2/DR* paradigms may simply depend on autophagy regulated by the ATG-16.2 WD40 domain to a lesser degree, or on a step of the autophagy pathway that the loss of the WD40 domain does not limit. Whichever applies, the ATG-16.2 WD40 domain is required for longevity induced by germline removal but not for the other paradigms tested, indicating that *glp-1/NotchR* animals depend on this domain in a way the others do not.

Loss of the ATG-16.2 WD40 domain altered three features of germline-less animals, each involving lysosomes. First, nuclear localization of the TFEB ortholog HLH-30, a transcription factor regulating lysosomal and autophagy genes [15], was reduced in intestinal cells. HLH-30 is itself required for longevity in *glp-1/NotchR* mutants [54], and thus, this partial regulation may explain how *glp-1/NotchR* mutants require the ATG-16.2 WD40 domain for their longevity. Our observation extends the reported requirement for the ATG16L1 WD40 domain in TFEB activation under lysosomal stress in mammalian cells [12–14] to a physiological, unstressed context. Second, elevated lysosomal pH in neurons of germline-less animals was abolished, and neuronal degradation of GFP::LGG-1 was reduced. Because neuronal lysosomal pH was unaffected by loss of the WD40 domain in an otherwise wild-type background, this WD40-dependent regulation of lysosomal pH in neurons appears to be engaged specifically by germline removal. The mechanism for this lysosomal pH regulation remains to be determined, but the direction of the effect matches a recent study in which mammalian epithelial cells lacking the ATG16L1 WD40 domain showed lysosomal hyperacidification, attributed to the domain inhibiting V-ATPase assembly through interaction with a V1 subunit [56]. Third, germline-less animals resisted heat-induced lysosomal membrane damage, and this resistance was lost in *atg-16.2(ΔWD40)* mutants. This result mirrors the requirement for the WD40 domain in limiting such damage in an otherwise wild-type background. These findings are interesting in light of prior work showing that germline removal uniquely remodels the lysosomal compartment, producing enlarged lysosomes [42], in contrast to the smaller, tubular lysosomes described in daf-2/InR and eat-2/DR animals [42,57]. We note that all three lysosome-related changes by the WD40 deletion in *glp-1/NotchR* mutants remain correlative: they establish that several lysosomal properties depend on the ATG-16.2 WD40 domain in the one paradigm where the domain is required for longevity, but not that any one of them is causal for lifespan. Also, these lysosomal changes cannot yet be ordered relative to one another, both because they were measured in different tissues and because they may reflect a single upstream defect or several parallel ones. Manipulating each property independently of the WD40 domain, for example by controlling lysosomal acidification, will be required to resolve these open questions.

In summary, our study establishes the ATG-16.2 WD40 domain is required *in vivo* for autophagic degradation, for limiting lysosomal membrane damage under heat stress, and in a paradigm-selective manner, for *C. elegans* longevity. It also links this domain, for the first time *in vivo*, to lysosomal pH regulation during aging. Thus, it shows that this domain, previously characterized largely for its non-canonical role in CASM, also contributes to canonical autophagy in an intact animal, with both functions converge on lysosomes. More broadly, our findings suggest that interventions targeting autophagy in age-related and other diseases may need to account for both canonical and non-canonical ATG16 functions, and for their shared impact on lysosomal homeostasis.

## METHODS

### C. elegans strains

Strains were cultured and maintained at 20 °C, under standard conditions. For experiments with temperature-sensitive *glp-1(e2144)* mutants, both *glp-1(e2144)* and wild-type (WT) N2 animals were allowed to hatch at 20 °C for 6 h and then moved to 25 °C until day 1 of adulthood. *Escherichia coli* OP50 was used as a food source during normal strains maintenance. See **Table S1** for strains generated and used for this study.

### Generation of atg-16.2(sq2) mutant

The *atg-16.2(sq2) allele*, which lacks the WD40 domain of ATG-16.2 (L224-R534), was generated by CRISPR-Cas9-mediated genome editing using the ribonucleoprotein (RNP) injection method[29–31]. Two guide RNAs (crRNA sequences: AGAACAGAUUGCAAAAGCUA; GUCAGAAGACUUGUACACUC) flanking the WD40-encoding region were designed using CRISPOR (https://crispor.gi.ucsc.edu/) and synthesized by Synthego. Guide RNAs were complexed with Cas9 Nuclease (QB3) prior to injection. A single-stranded oligonucleotide (ssODN) repair template (GCTTTGCAAGAAGAACGCGCACAGATGCGAATCAGAGAACAGATTGCAAAAGCTACAGCTGATTTAAATTT AGGTGACGAGCGAGCATCTTCAGCATTTGGTACATCTCCCGAAACTTGATCTTCAATGTATGAAATTGAATATAATTGTACAG) was used to direct deletion of the WD40 domain-encoding region and to introduce silent mutations disrupting the PAM site. The *rol-6(su1006)* plasmid was used as a co-injection marker to enrich for edited animals. Roller F1 progeny were singled, and deletion alleles were identified by PCR and confirmed by Sanger sequencing. The *atg-16.2(sq2)* allele was outcrossed four times against the N2 wild-type strain before use in experiments.

### Construction of transgenic strains

The *rgef-1p::GFP::LGG-1(G116AG117A)* and *lgg-1p::GFP::LGG-1(G116AG117A)* vectors were made by site-directed mutagenesis using primers TGTATGCCGCAGAGGTCGAAAAGAAGGAATAACAAC and CCTCTGCGGCATACACACTTTCGTCACTGTAGGC. To construct pMH1545 *rgef-1p::GFP::LGG-1(G116A G117A)*, pMH1264 (*rgef-1p::GFP::LGG-1(G116A)* [38]) was used to change glycine 117 (GGA) to alanine 117 (GCA) by site-directed mutagenesis. To construct pMH1546 *lgg-1p::GFP::LGG-1(G116A G117A)*, pMH1298 (*lgg-1p::GFP::LGG-1(G116A)*) was used to change glycine 117 (GGA) to alanine 117 (GCA) by site-directed mutagenesis. The final products (pMH1545 and pMH1546) were verified by sequencing. Plasmid DNA was prepared using the Miniprep kit (Qiagen). Transgenic animals expressing an extrachromosomal array were created by gonadal microinjection of pMH1545 or pMH1546 with the indicated co-injection marker into the strains indicated in Table S1, generating MAH1479 and MAH1491, respectively.

### Immunoblotting

For endogenous LGG-1 lipidation assays, animals were collected in M9 buffer, washed twice to remove bacteria, and resuspended in 18 μL of M9 buffer. 6 μL of 4x Laemmli Sample Buffer (Bio-Rad, #1610747) supplemented with 10% 2-Mercaptoethanol (Bio-Rad, #1610710) was added, and the resuspension were snap-frozen in liquid nitrogen and thawed twice. For ATG-16.2::mScarlet detection and lysosomal degradation assays, animals were collected in S-basal buffer (100 mM NaCl, 50 mM potassium phosphate, 5 μg/mL cholesterol, pH 6.0), washed twice to remove bacteria, and snap-frozen in liquid nitrogen. Frozen pellets were resuspended in 50 μL of lysis buffer (100 mM Tris, pH 8.0, 150 mM NaCl, 1% Triton X-100, 1% deoxycholic acid, 0.1% SDS, 5 mM EDTA, and 10 mM NaF) [42] supplemented with Halt Protease Inhibitor Cocktail (Thermo Scientific, #1861279) and were sonicated using Bioruptor 300 (Diagenode) by 10 cycles (30 sec sonication, 30 sec rest) at high power. Lysates were cleared by centrifugation at 3,000 x g, 4 min, 4°C, and 10 μL of Laemli SDS sample buffer, reducing (6X) (Thermo Scientific, #J61337) were added to 50 μL of the lysates. Proteins were separated on SDS-PAGE gels and transferred to PVDF membranes. Membranes were blocked in 5% skim milk in TBST and incubated overnight at 4°C with primary antibodies against anti-LGG-1 (Abgent, 1:1,000 dilution), HRP-conjugated anti-GFP (Abcam, ab190584, 1:1000 dilution), anti-RFP (Proteintech, 6G6, 1:1,000 dilution), and anti-β-Actin (Cell Signaling, 3700S, 1:2,500 dilution), followed by HRP-conjugated anti-mouse IgG (Cell Signaling, 7074S, 1:2,000 dilution) and anti-rabbit IgG (Cell Signaling, 7076S, 1:2,000 dilution). Signal was detected with Pierce ECL Western Blotting Substrate (Thermo, #32106), SuperSignal West Pico PLUS Chemiluminescent Substrate (Thermo Scientific, #34577), SuperSignal West Dura Extended Duration Substrate (Thermo Scientific, #34076), and SuperSignal West Femto Maximum Sensitivity Substrate (Thermo Scientific, #34095) on Azure 600 (Azure Biosystems). Band intensities were quantified in FIJI [58].

Protein expression of ATG-16.2(±WD40)::mScarlet was assessed on day 1 of adulthood in WT, *atg-16.2(ΔWD40)*, *glp-1(e2144)*, and *atg-16.2(ΔWD40); glp-1(e2144)* animals (MAH1997, MAH1961, MAH1975 and MAH1996, respectively, see Table S1). Approximately 100 animals per genotype were collected and processed for immunoblotting as described above. ATG-16.2(±WD40)::mScarlet was resolved on 4–12% gradient gels and detected with an anti-RFP antibody. Welch’s t-test or one-way ANOVA followed by Tukey’s multiple comparisons was applied to the means from biological replicates using GraphPad Prism.

Lipidation of endogenous LGG-1 was assessed on day 1 of adulthood in WT, *atg-5(lof)* and *atg-16.2(ΔWD40)* animals (N2, MAH1389 and MAH1354, respectively, see Table S1). Approximately 100 animals per genotype were collected and processed for immunoblotting as described above. Unlipidated LGG-1-I and lipidated LGG-1-II were resolved on 12% gels and detected with an anti-LGG-1 antibody. The LGG-1-II/LGG-1-I ratio was calculated for each genotype and normalized to WT within each biological replicate. One-way ANOVA followed by Tukey’s multiple comparisons was applied to the means from biological replicates using GraphPad Prism.

Lysosomal degradation of autophagic cargo was assessed using the neuron-specific *rgef-1p::gfp::lgg-1* reporter as described previously (Wang et al., 2025), in WT, *atg-16.2(ΔWD40)*, *glp-1(e2144)*, and *atg-16.2(ΔWD40); glp-1(e2144)* animals (MAH242, MAH1358, MAH250 and MAH1928, respectively, see Table S1). Approximately 100 animals per genotype were collected on day 3 of adulthood and processed for immunoblotting as described above. Unlipidated GFP::LGG-1-I, lipidated GFP::LGG-1-II and free GFP were resolved on 12% gels and detected with an HRF-conjugated anti-GFP. Lipidation was calculated as the GFP::LGG-1-II/GFP::LGG-1-I ratio and degradation as free GFP relative to total GFP signal (free GFP plus GFP::LGG-1-I and GFP::LGG-1-II). Welch’s t-test or one-way ANOVA followed by Tukey’s multiple comparisons was applied to the means from biological replicates using GraphPad Prism.

### Healthspan assays

Thrashing rate was measured by counting the body bends of 6 animals transferred to M9 medium in 24-well plates for 30 seconds. One-way ANOVA followed by Tukey’s multiple comparisons was applied to the means from biological replicates using GraphPad Prism.

Pharyngeal pumping was measured by counting the grinder movements in the pharyngeal bulb for 30 seconds. One-way ANOVA followed by Tukey’s multiple comparisons was applied to the means from biological replicates using GraphPad Prism.

For assessment of reproductive output, ∼10 animals were singled on 6 cm NGM plates at the L4 larval stage then transferred daily onto fresh plates during the reproductive span up to day 5 of adulthood. The number of larvae produced by each animal per day was counted then summed up. One-way ANOVA followed by Tukey’s multiple comparisons was applied to the means from biological replicates using GraphPad Prism.

On day 10 of adulthood, ∼30 animals were removed from the NGM plates and suspended for 3 h in liquid cultures of standard OP50 bacteria (grown overnight) mixed with blue food dye (Spectrum FD&C Blue #1 PD110, 5.0% wt/vol in water). Animals were then transferred to NGM plates seeded with OP50 bacteria and analyzed for the presence or absence of blue food dye in the body cavity using a Leica DFC310 FX microscope under 40x magnification. Animals with dye present only in the germline were excluded from the analysis. Logistic regression with Tukey’s multiple correction was applied to the means from biological replicates using RStudio.

### Autophagy measurement

Autophagy was assessed by quantifying SQST-1/p62::GFP fluorescence intensity in the head region, or GFP::LGG-1/Atg8 puncta in nerve-ring neurons, proximal intestinal cells and body-wall muscle.

For SQST-1/p62::GFP imaging[27], fluorescence intensity of SQST-1::GFP was measured in the head region on day 1 adults (MAH349, MAH1198 and MAH1423, see **Table S1**). Animals were imaged on empty NGM plates after anesthetization with M9 medium containing 0.1% sodium azide. Images were acquired with a Leica DFC310 FX camera at 500 ms of exposure. The mean fluorescence intensity was measured in the head region by outlining the same-sized circular region of interest using FIJI software[58] per animal, normalized to control animals. One-way ANOVA followed by Tukey’s multiple comparisons was applied to the means from biological replicates using GraphPad Prism.

For autophagy flux assays of GFP::LGG-1/Atg8 puncta[27], approximately 10 animals of day 1 adults were cultured on OP50-seeded 35 mm NGM plates containing DMSO (0.01% for *rgef-1p::gfp::lgg-1* reporter and 0.16% for *lgg-1p::gfp::lgg-1* reporter) or BafA (2.5 μM for *rgef-1p::gfp::lgg-1* reporter and 40 μM for *lgg-1p::gfp::lgg-1* reporter) for 1 (*lgg-1p::gfp::lgg-1*) or 2 (*rgef-1p::gfp::lgg-1* reporter) hours. Strains used were: MAH242 and MAH1358 for *rgef-1p::gfp::lgg-1*, DA2123 and MAH1388 for *lgg-1p::gfp::lgg-1* (see **Table S1**). Animals were raised at 20 °C. For imaging and puncta quantification, animals were mounted on a 3% agarose pad in M9 medium containing 0.1% NaN_3_ for imaging using Zeiss Imager Z2 with a Zeiss Apotome 3 optical sectioning unit, a Zeiss Axiocam 712 mono camera, and Zen 3.4 software. Number of GFP::LGG-1/Atg8 positive puncta in body-wall muscle, two proximal intestinal cells were counted at 100X magnification. For nerve-ring neurons, Z-stack images were taken at 1 μm slice intervals for puncta quantification and were used to generate max Z-projection images for quantification. Two-way ANOVA followed by Tukey’s multiple comparisons was applied to the means from biological replicates using GraphPad Prism.

Lipidation-deficient GFP::LGG-1(G116AG117A) reporters (MAH1749 and MAH1506 for *rgef-1p::gfp::lgg-1(G116AG117A)*, and MAH1491 and MAH1966 for *lgg-1p::gfp::lgg-1(G116AG117A)*; see Table S1) were imaged and quantified identically, under basal conditions only. Welch’s t-test was applied to the means from biological replicates.

### Aggregate counting

The number of neuronal Q40::YFP aggregates was quantified in individual on day 7 of adulthood (MAH1308, MAH1475 and MAH1477, see **Table S1**). The number of muscle Q35::YFP aggregates was quantified in individual on day 4 of adulthood (AM140, MAH1474 and MAH1485, see **Table S1**). Animals were analyzed on NGM plates without food after anaesthetization with M9 medium containing 0.1% NaN_3_. Representative images were acquired by mounting animals on a 3% agarose pad in M9 medium containing 0.1% NaN3 and using Zeiss Imager Z2 with a Zeiss Apotome 3 optical sectioning unit, a Zeiss Axiocam 712 mono camera and Zen 3.4 software. One-way ANOVA followed by Tukey’s multiple comparisons was applied to the means from biological replicates using GraphPad Prism.

### Lysosomal pH measurement

Lysosomal pH was assessed by quantifying mTFP/mCherry fluorescence intensity ratio of neuronal FIRE-pHLy (mTFP::LMP-1::mCherry) strains MAH1497, MAH1742, MAH1542 and MAH1741 (see Table S1). Animals on day 3 adulthood were mounted on a 3% agarose pad in M9 medium containing 0.1% NaN_3_ for imaging using Zeiss Imager Z2 with a Zeiss Axiocam 712 mono camera and Zen 3.4 software. 40X magnification. Z-stack images were taken at 1 μm slice intervals and were used to generate max Z-projection images for quantification. To quantify FIRE-pHLy fluorescence, mCherry was used as the reference channel for segmenting lysosomes as described previously[43]. Object masks for lysosomes from mCherry images were directly applied to 488 nm source images to segment mTFP identically as mCherry. The mean fluorescence intensities of mCherry and mTFP channels were generated, and the ratios were calculated. All measures were outputted as a Microsoft Excel file for further analysis. Welch’s t-test or one-way ANOVA followed by Tukey’s multiple comparisons was applied to the means from biological replicates using GraphPad Prism.

### Lysosomal damage assay

Lysosomal membrane damage was assessed using a ubiquitously expressed mAG::hLGALS3 reporter (*eft-3p::mAG::hLGALS3*) [45]. Strains used were AGD3761, MAH1727, MAH1746 and MAH1807 (see Table S1). On day 3 of adulthood, animals were exposed to 36°C for 3 h and allowed to recover at 20°C for 3 h before imaging; control animals were maintained at 20°C throughout. Animals were imaged on empty NGM plates after anesthetization with M9 medium containing 0.1% sodium azide. Images were acquired with a Leica DFC310 FX camera at 500 ms of exposure. mAG::hLGALS3 puncta were quantified in the whole-body by manual counting. Two-way ANOVA followed by Tukey’s multiple comparisons was applied to the means from biological replicates using GraphPad Prism.

### Thermotolerance assay

Thermotolerance was assessed on day 3 of adulthood. Animals were raised at 20°C, except for *glp-1(e2144)*, which were shifted as described under *C. elegans strains*. Approximately 20 animals per genotype were transferred to 4–5 plates and exposed to 36°C in incubator for the durations indicated in the figure legends (5 h for *eat-2(ad1116)*, 9 h for *daf-2(e1370)* and 6 h for *glp-1(e2144)*). Exposure times were selected per paradigm to produce scoreable mortality, so comparisons are made within and not across paradigms. After heat shock, animals were returned to 20°C and survival was scored 24 h (*eat-2, glp-1*) or 48 h (*daf-2*) later; animals were scored as dead if they failed to respond to gentle touch with a platinum wire. Logistic regression with Tukey’s multiple comparisons was applied to the means from biological replicates using RStudio.

### Lifespan analysis

Lifespan was measured at 20 °C using six 6 cm NGM or NGM/Carb plates seeded with OP50 bacteria or dsRNA-expressing bacteria with 15–20 animals per plate. The L4 larval stage was recorded as day 0 of the lifespan and animals were transferred at least every other day to new 6 cm NGM plates throughout the reproductive period. Censoring occurred if animals desiccated on the edge of the plate, escaped, ruptured or suffered from internal hatching. Statistical analysis by log-rank test was performed using Oasis 2[59]. *P*-values were calculated with the log-rank (Mantel–Cox) method. See Supplementary Tables for a summary of all lifespan experiments.

### HLH-30::GFP nuclear localization

Nuclear localization of HLH-30::GFP was scored as described previously [54] on day 3 of adulthood, in strains MAH. . Animals were imaged on empty NGM plates after anesthetization with M9 medium containing 0.1% sodium azide. Images were acquired with a Leica DFC310 FX camera at 500 ms of exposure. and each animal was classified as nuclear or non-nuclear manually. Logistic regression with Tukey’s multiple comparisons was applied to the means from biological replicates using RStudio.

### Statistical analysis

GraphPad Prism (version 11.0.2) and RStudio (version 2026.05.0-208) were used for statistical analysis. Comparisons with p < 0.05 were considered statistically significant except for lifespan analysis where comparisons with p < 0.01 were considered statistically significant.

---

**Figure S1:**
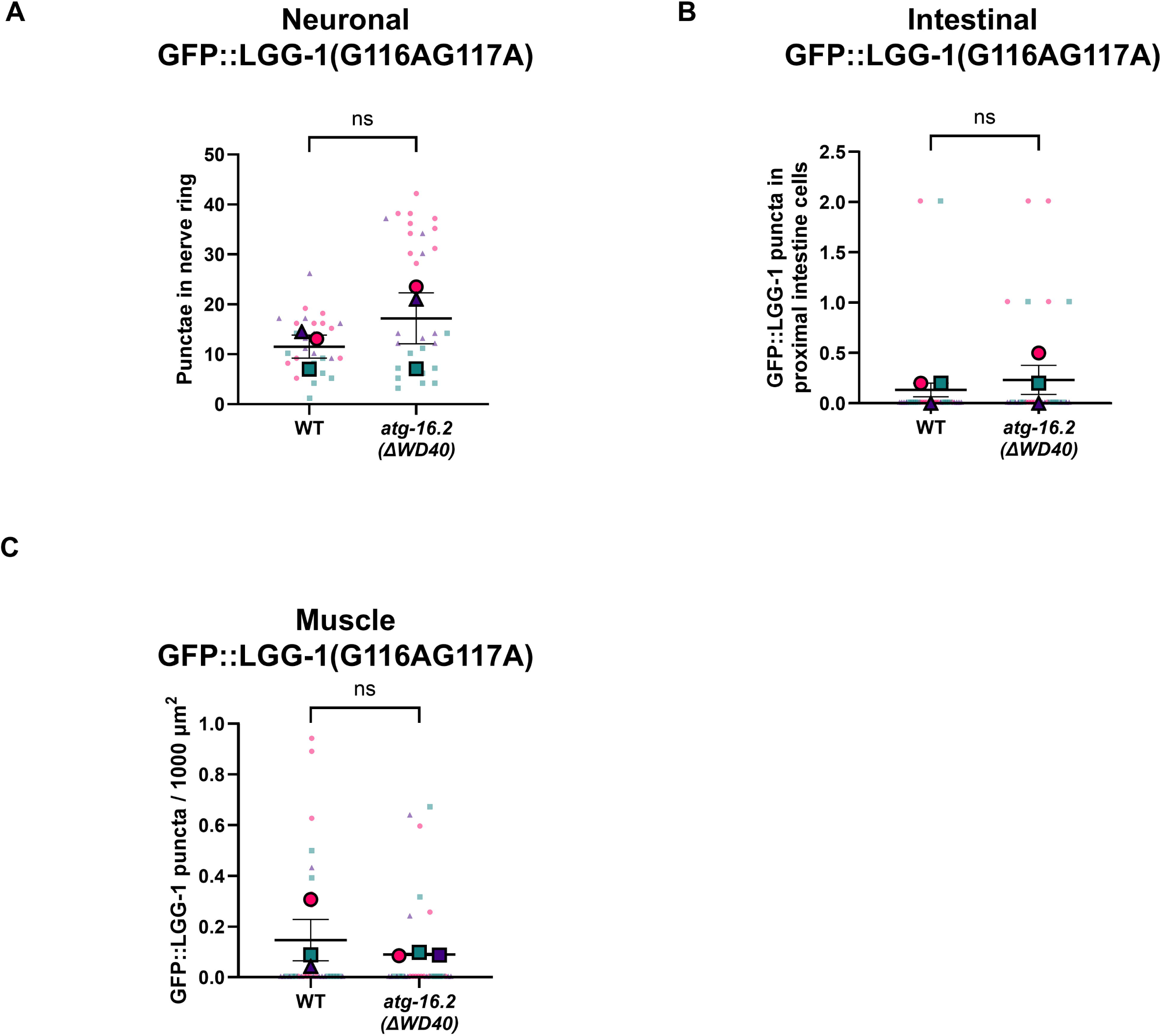
Lipidation-deficient GFP::LGG-1(G116AG117A) forms fewer puncta relative to GFP::LGG-1, and its puncta numbers are unaffected by loss of the ATG-16.2 WD40 domain. **A.** Quantification of GFP::LGG-1(G116AG117A) puncta in the nerve rings of WT and *atg-16.2(ΔWD40)* animals. Small dots represent data from each animal and large dots represent the means from independent experiments. Error bars show the mean ± SEM. n = 3. 30 animals. Welch’s t-test. **B.** Quantification of GFP::LGG-1(G116AG117A) puncta in proximal intestinal cells of WT and *atg-16.2(ΔWD40)* animals. Small dots represent data from each animal and large dots represent the means from independent experiments. Error bars show the mean ± SEM. n = 3. 30 animals. Welch’s t-test. **C.** Quantification of GFP::LGG-1(G116AG117A) puncta in body wall muscles of WT and *atg-16.2(ΔWD40)* animals. Small dots represent data from each animal and large dots represent the means from independent experiments. Error bars show the mean ± SEM. n = 3. 30 animals. Welch’s t-test.

**Figure S2:**
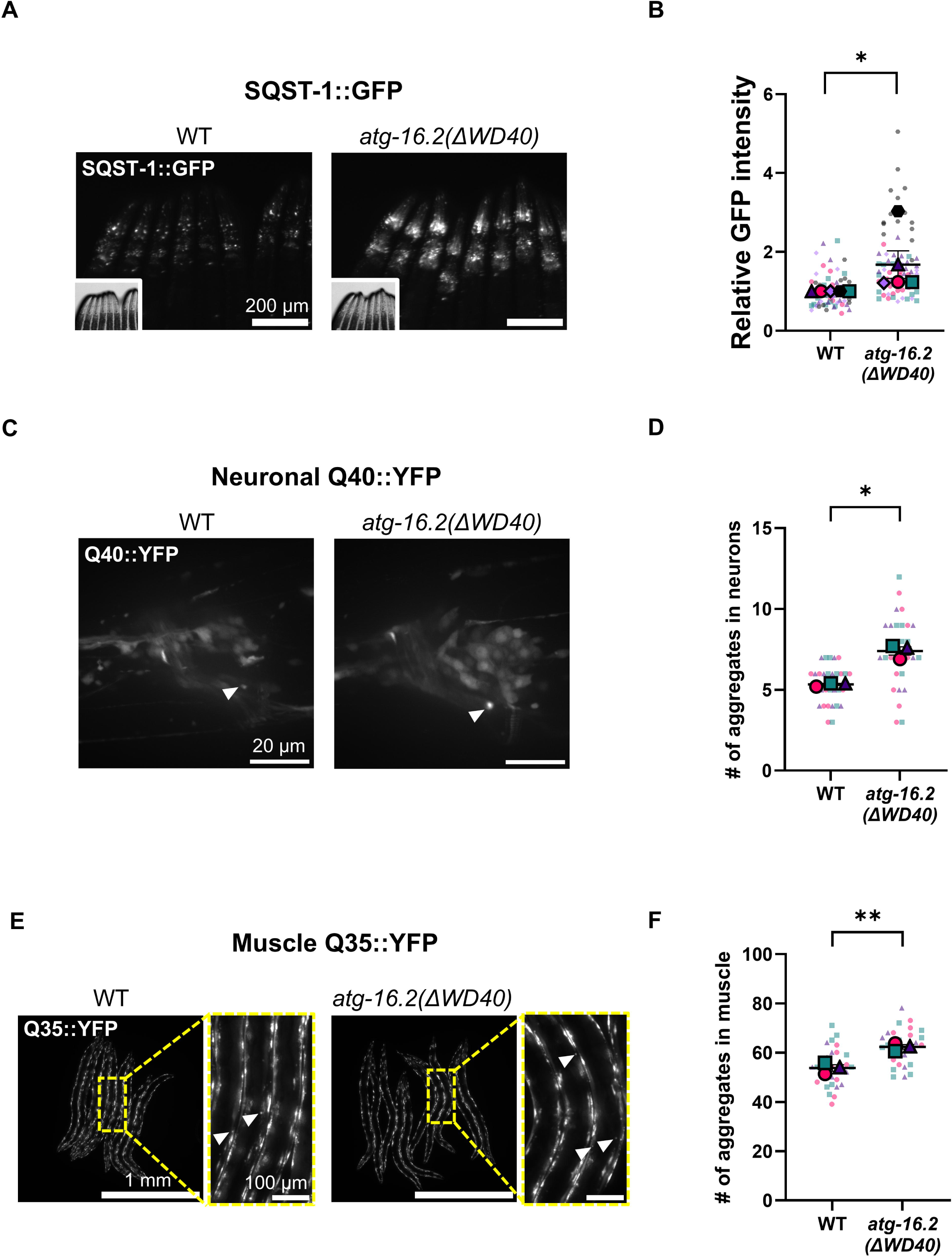
*atg-16.2(ΔWD40)* mutation leads to accumulation of the autophagy substrates SQST-1/p62 and polyQ aggregates. **A.** Representative images of SQST-1/p62::GFP reporter strains carrying WT or *atg-16.2(ΔWD40)* mutation on day 1 of adulthood. Scale bar: 200 μm. **B.** Quantification of SQST-1/p62::GFP fluorescence intensity shown in (A). Small dots represent data from each animal and large dots represent the means from independent experiments. Error bars show the mean ± SEM. n = 5. 78 animals. Welch’s t-test. Statistical tests were performed on raw measurements; values are displayed normalized to the WT control mean within each independent experiment for presentation. **C.** Representative images of neuronal Q40::YFP reporter strains carrying WT or *atg-16.2(ΔWD40)* mutation on day 7 of adulthood. Arrowheads indicate representative puncta. Scale bar: 20 μm. **D.** Quantification of Q40::YFP puncta in the head region. Small dots represent data from each animal and large dots represent the means from independent experiments. Error bars show the mean ± SEM. n = 3. 30 animals. Welch’s t-test. *:p<0.05. **E.** Representative images of muscle Q35::YFP reporter strains carrying WT or *atg-16.2(ΔWD40)* mutation on day 4 of adulthood. Arrowheads indicate representative puncta. Scale bar: 1 mm in the large pictures and 100 μm in the zoom-in images. **F.** Quantification of Q35::YFP puncta in body wall muscle. Small dots represent data from each animal and large dots represent the means from independent experiments. Error bars show the mean ± SEM. n = 3. 30 animals. Welch’s t-test. **:p<0.01.

**Figure S3:**
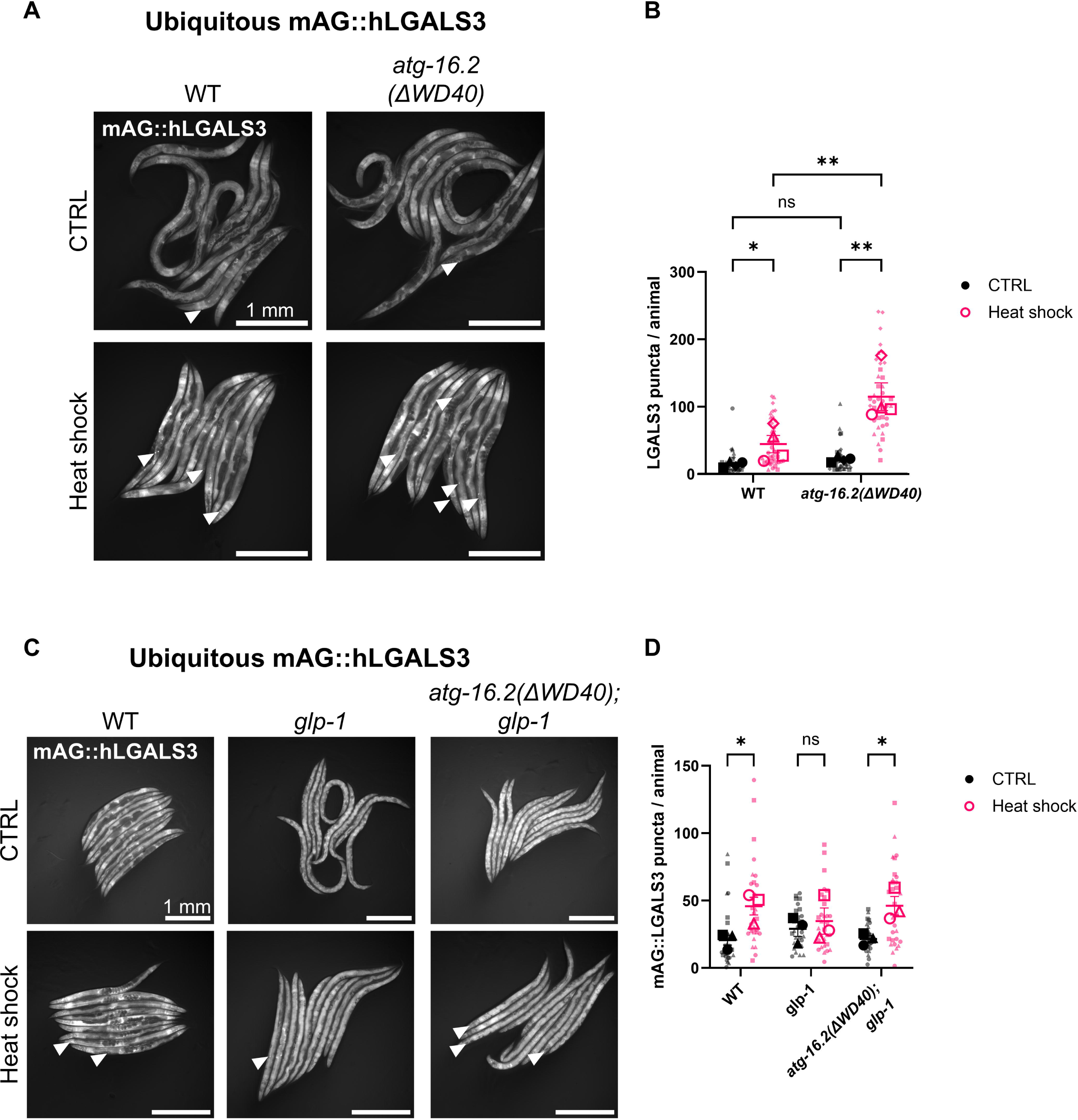
*atg-16.2(ΔWD40)* mutation makes lysosomes more susceptible to damage after heat shock in otherwise wild-type animals and *glp-1* mutants. **A.** Representative images of mAG::hLGALS3 reporter animals in WT and *atg-16.2(ΔWD40)* backgrounds under control conditions (CTRL) and after heat shock (HS; 36 °C for 3 h followed by 3 h of recovery at 20 °C) on day 3 of adulthood. Arrowheads indicate representative puncta. Scale bar: 1 mm. **B.** Quantification of mAG::hLGALS3 puncta in the whole body shown in (A). Small dots represent data from each animal and large dots represent the means from independent experiments. Error bars show the mean ± SEM. n = 4. 40 animals. Two-way ANOVA with Fisher’s LSD test. *: p<0.05, **: p<0.01. **C.** Representative images of mAG::hLGALS3 reporter animals in WT, *glp-1(e2144)* and *atg-16.2(ΔWD40); glp-1(e2144)* backgrounds under control conditions (CTRL) and after heat shock (HS; 36 °C for 3 h followed by 3 h of recovery at 20 °C) on day 3 of adulthood. Arrowheads indicate representative puncta. Scale bar: 1 mm. **D.** Quantification of mAG::hLGALS3 puncta shown in the whole body shown in (C). Small dots represent data from each animal and large dots represent the means from independent experiments. Error bars show the mean ± SEM. n = 3. 30 animals. Two-way ANOVA with Tukey’s multiple comparisons. *: p<0.05.

**Figure S4:**
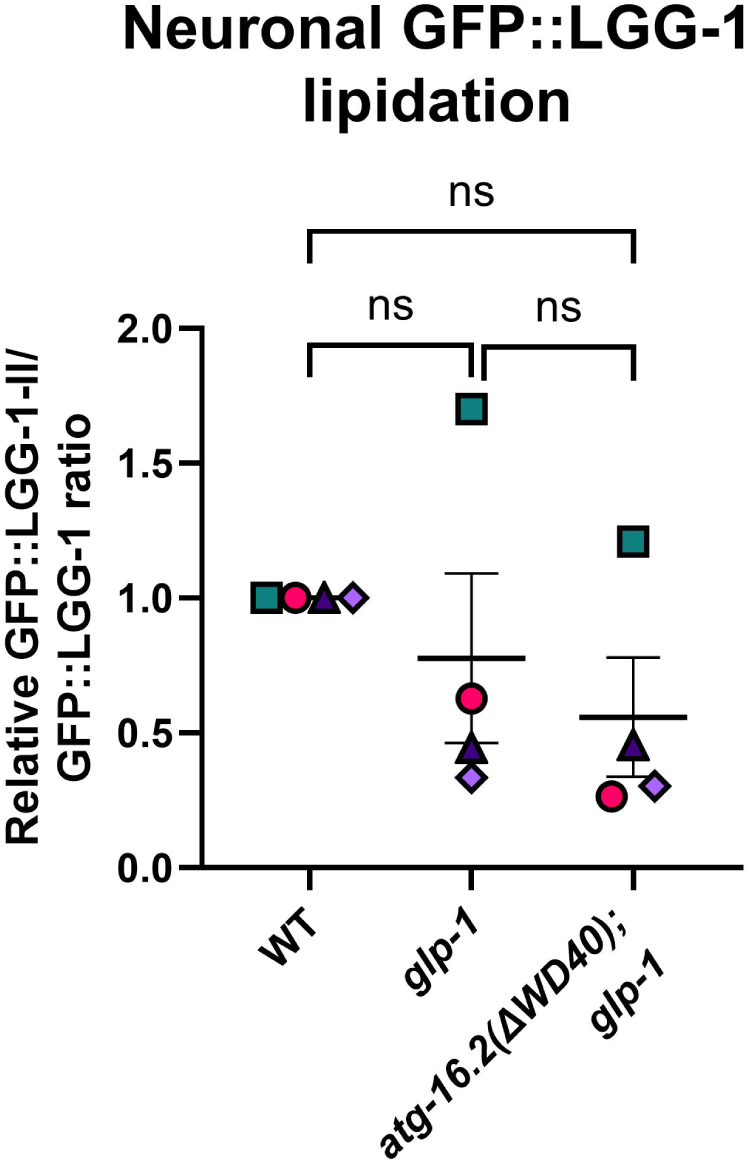
*atg-16.2(ΔWD40)* mutation does not affect LGG-1 lipidation in neurons of *glp-1* mutants. Quantification of lipidation from Figure 5D, calculated as the GFP::LGG-1-II/GFP::LGG-1-I ratio and normalized to WT within each replicate. Error bars show the mean ± SEM. n = 4. One-way ANOVA with Tukey’s multiple comparisons. Statistical tests were performed on log2 values of raw measurements; values are displayed normalized to the WT control mean within each independent experiment for presentation.

## ACKNOWLEDGEMENTS

We thank Meagan Arnold from Monica Driscoll’s lab for help designing CRISPR sgRNAs (CRISPR-design) and Maria Sallee from M.H.’s lab for helping generating GFP::LGG-1(G116AG117A) transgenic animals. Some of the nematode strains used in this work were provided by the *Caenorhabditis* Genetics Center (University of Minnesota), which is supported by the NIH Office of Research Infrastructure Program (P40 OD010440). We acknowledge WormBase WS283. This work was funded by the Paul F. Glenn Center for Biology of Aging at the Buck Institute for Research on Aging, Larry L. Hillblom Fellowship 2023-A-019-FEL and a Valley Foundation Fellowship to H.E., a Taiwan Global Fellowship to L.-H.S., NIH/NIA grant R01NS127414 and RF1NS142354 to A.W.K, and NIH/NIA grant R01AG038664 to M.H. The content of this article is solely the responsibility of the authors and does not necessarily represent the official views of the NIH.

## AUTHOR CONTRIBUTIONS

A. E. and M. Hansen designed research; H.E., L.H.S, S.K.D, D.B., S.D., and A.L. performed research; M. Hodul and A.W.K contributed new reagents/analytic tools; H.E., L.H.S, and S.K.D. analyzed data; and H.E. wrote the original manuscript with input from M. Hansen, and other co-authors provided feedback on the final version.

## Declaration of AI and AI-assisted technologies

Claude.ai was used in the writing process for minor grammatical refinement during the drafting of this article.

---

**Table S1:**
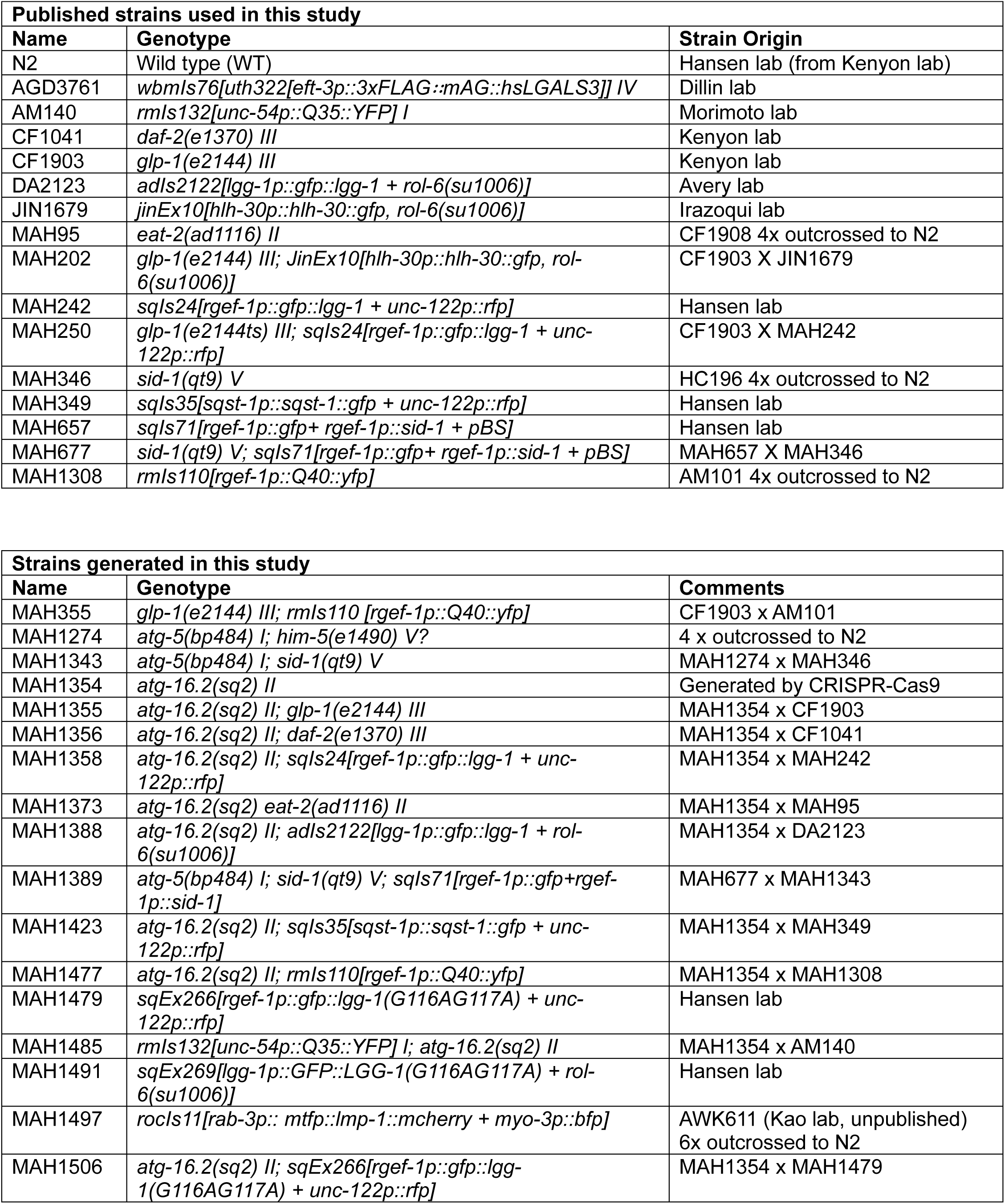
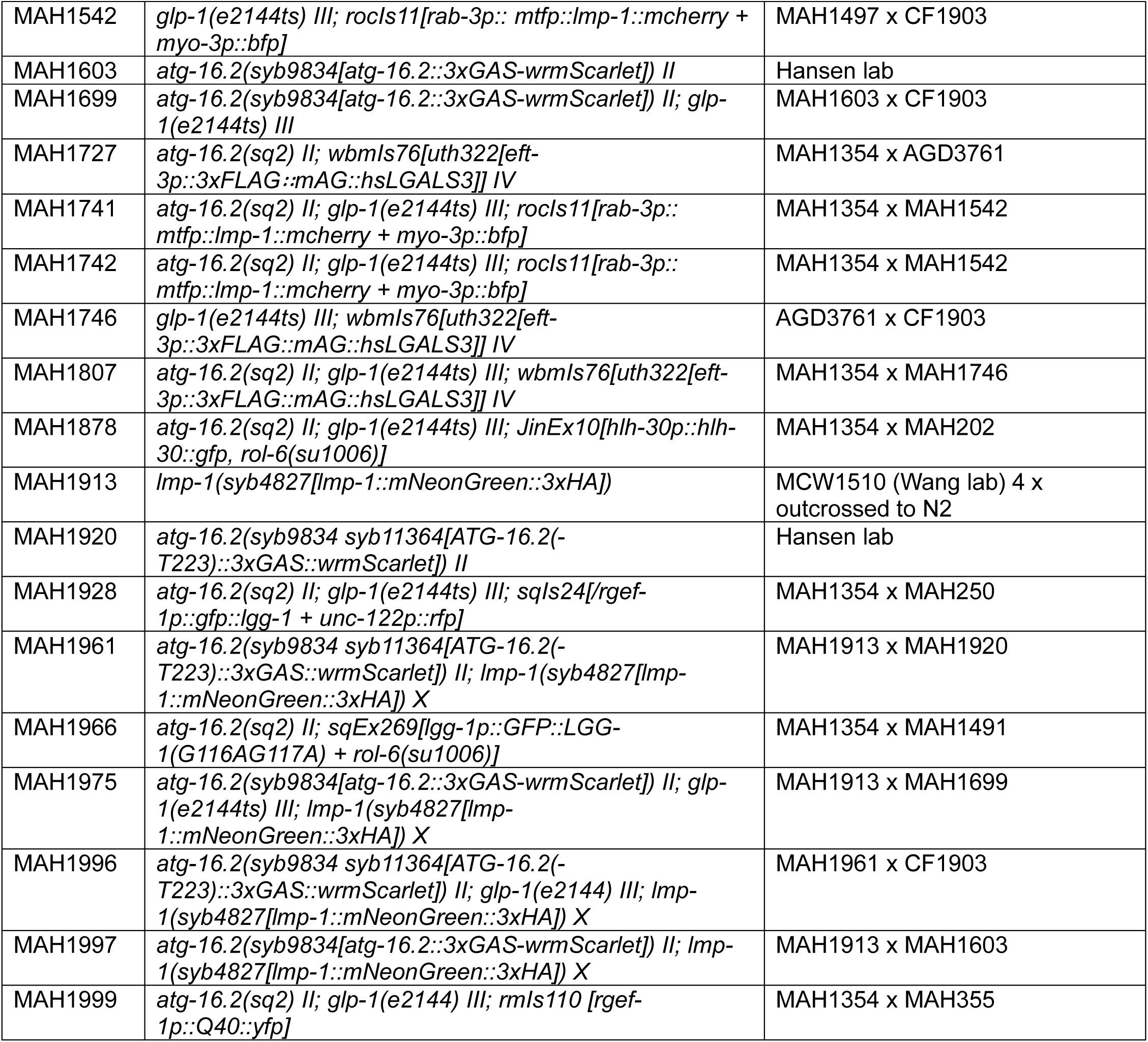
Strains used in this study.

